# Understanding cognitive function in post-COVID-19 condition with fatigue: an interdisciplinary multimodal investigation

**DOI:** 10.64898/2026.10.21.760664

**Authors:** Ulrika Birberg Thornberg, Anders Eklund, Kristin Zeiler, Sofia Morberg Jämterud, Felipe León, Evren Özarslan, Agnes Andersson, Richard Levi, David Engblom, Anestis Divanoglou, Ida Blystad

## Abstract

Lingering cognitive symptoms of post-COVID-19 condition (PCC) continue to challenge our understanding, highlighting the need for interdisciplinary research. We investigated whether individuals with PCC and fatigue differ from matched controls across neurocognitive, neuroimaging, and immunological measures, integrating these findings with qualitative phenomenological analyses through multimodal cross-analysis.

Individuals with PCC and clinically significant fatigue (n=34), and age-, sex-, and education-matched controls recovered from COVID-19 without PCC (n=37) were included. Assessments encompassed physician evaluation, neuropsychological and functional tests, structural and functional MRI, immunological tests, and qualitative phenomenological analysis of interviews. Data were analyzed using established methods and an interdisciplinary cross-analysis to examine persistent cognitive symptoms in PCC.

Group comparisons revealed subtle differences between individuals with PCC and controls, while neurocognitive performance and immunological markers remained within normative ranges. Modest differences in white-matter, myelin-related, and global brain measures were observed, alongside differences in associations between network connectivity and neurocognitive performance across groups, suggesting reduced network resilience rather than domain specific deficits. This pattern resonated with qualitative phenomenological findings: altered experiences of concentrating and difficulties to execute decisions and actions, despite preserved performance on standard neurocognitive tests.

These findings suggest that cognitive symptoms in post-COVID-19 fatigue are associated with subtle neurobiological alterations not captured by standard cognitive assessments. In this highly fatigued sample, converging neuroimaging, neurocognitive and immunological findings point toward reduced network efficiency rather than overt cognitive impairment. Qualitative phenomenological analysis further highlighted altered modes of being in the world characterized by increased cognitive effort and difficulties executing decisions and actions.

## 1. Introduction

Post-COVID-19 Condition (PCC) occurs after SARS-CoV-2 infection, typically with symptoms emerging within three months and persisting for at least two months. Increasing evidence characterizes PCC as a heterogeneous but clinically significant condition that impairs daily functioning and social participation.^1,2^ Common symptoms include fatigue, cognitive deficits, sleep disturbances, depression, and anxiety. Among these, fatigue and cognitive deficits are the most frequent and disabling neuropsychiatric features.^3,4^ Meta-analytic data indicate that fatigue affects approximately 43% of individuals with PCC, while cognitive impairment, memory deficits and sleep disturbances occur in 24–28%.^5^

Self-reported cognitive difficulties, including problems with memory, attention, and executive function, show considerable heterogeneity and do not always align with findings from functional MRI, EEG, comprehensive neurocognitive assessment, or immunological assessments.^4,6^ This mismatch highlights the complexity of characterizing cognitive dysfunction in PCC and underscores the need for multimodal approaches integrating ‘subjective’ and ‘objective’ measures. Recent reflections on the “subjectivity–objectivity gap,” have highlighted how divergences between reported and measured cognitive difficulties may complicate clinical evaluation,^7^ while also problematizing the idea of a clear-cut distinction between objective and subjective dimensions.^8^

Brain alterations have been documented during acute SARS-CoV-2 infection and in PCC.^9^ Meta-analytic evidence suggests that approximately 43% of patients undergoing CT/MRI show neuroimaging abnormalities, most commonly white-matter changes and non-specific ischemic findings.^9^ A review integrating structural MRI, diffusion imaging, fMRI, and perfusion imaging across 1,219 individuals with PCC reported convergent evidence of reduced gray-matter volume and cortical thickness, increased diffusivity, decreased fractional anisotropy, white-matter tract injury, disrupted network connectivity, and reduced cerebral blood flow.^10^ Even though these alterations may underlie neurological and psychiatric symptoms in PCC, their relationship to measured cognitive deficits remains unclear, with evidence remaining heterogeneous. Several multimodal MRI studies have linked brain alterations to measurable cognitive deficits after COVID-19, including deficits in memory, processing speed and executive function.^11–13^

While some studies report correlations between neuroimaging markers and objective neurocognitive deficits, others report neural alterations without corresponding cognitive associations,^14^ indicating variability across cohorts, methods, and cognitive assessments. This heterogeneity is also reflected in reviews summarizing inconsistent structural and functional MRI findings across PCC cohorts.^15^

Despite a growing number of neuroimaging studies in PCC, relatively few combine advanced MRI with comprehensive neuropsychological testing and immunology markers. Even fewer include well-matched control groups, and some lack controls altogether, limiting interpretability of brain–behavior relationships.^2^ Consequently, the link between neurobiological alterations and cognitive impairment in PCC remains unclear, underscoring the need for studies integrating detailed cognitive testing with multimodal MRI in controlled designs.

Qualitative studies consistently identify cognitive symptoms as central and life-altering in PCC,^16^ yet multimodal studies have not incorporated patients’ lived experience perspectives. Patients report fluctuating impairments in attention, memory, executive function, and language that restrict everyday functioning, work ability, and social interaction, and are intertwined closely with fatigue.^16,17^ These difficulties are not only cognitively disabling but also psychosocially disruptive, contributing to a loss of identity, disruption of everyday roles, and anxiety about the future.^16^ Together, qualitative evidence suggests that cognitive symptoms are experienced as embodied, relational, and existential, reshaping how individuals live and understand themselves after Covid-19. The current study took inspiration from an enactive approach that acknowledges embodied subjectivity and the living organism as two dimensions of the living being as a person, embedded in an environment.^18^ This approach allows for a holistic focus, which is apt for the present study (see ^19^).

The aim of this study was to investigate whether individuals with PCC and fatigue differ from matched controls across structural and functional MRI, neurocognitive performance, and immunological (cytokine) profiles, and whether these differences relate to findings from qualitative phenomenological analysis of lived experience. We further examined associations across modalities.

## 2. Methods

### 2.1. Study design and population

This interdisciplinary study employs a cross-sectional case–control design combining neuroradiological, neurocognitive, immunological assessments, and a qualitative phenomenological analysis. It includes 71 individuals: 34 with PCC and 37 age-, sex-, and education-matched controls who had recovered from COVID-19 without developing PCC. Participants were recruited in Östergötland, Sweden, between October 2022 and June 2024.

### 2.2. Procedure

Eligible participants were Swedish-speaking adults aged 20–65 years with confirmed SARS-CoV-2 infection at least six months before inclusion. Participants with PCC were recruited from a previous COVID-19 cohort whose members had consented to future contact, and through information posted at regional PCC clinics. Controls were recruited through announcements at hospitals, the university, and social media. Interested individuals provided written informed consent before eligibility screening by structured telephone interview. Individuals with PCC and clinically significant fatigue (Multidimensional Fatigue Inventory, MFI ≥ 53) were compared with matched controls without PCC and minimal fatigue (MFI < 53). Exclusion criteria included pre-existing or comorbid conditions that could affect cognition or explain fatigue, as well as sensory or motor impairments precluding valid neuropsychological testing, and MRI contraindications.

Participants meeting inclusion criteria who provided informed consent underwent blood and urine sampling, followed by a comprehensive clinical assessment, comprising neuropsychological, medical, and physiotherapy evaluations. Brain MRI was performed separately within a few weeks and evaluated by a neuroradiologist. All participants with PCC additionally completed a phenomenological interview. Additional methodological details are available in the study protocol.^19^ Figure 1 shows participant flow.

**Figure 1.**
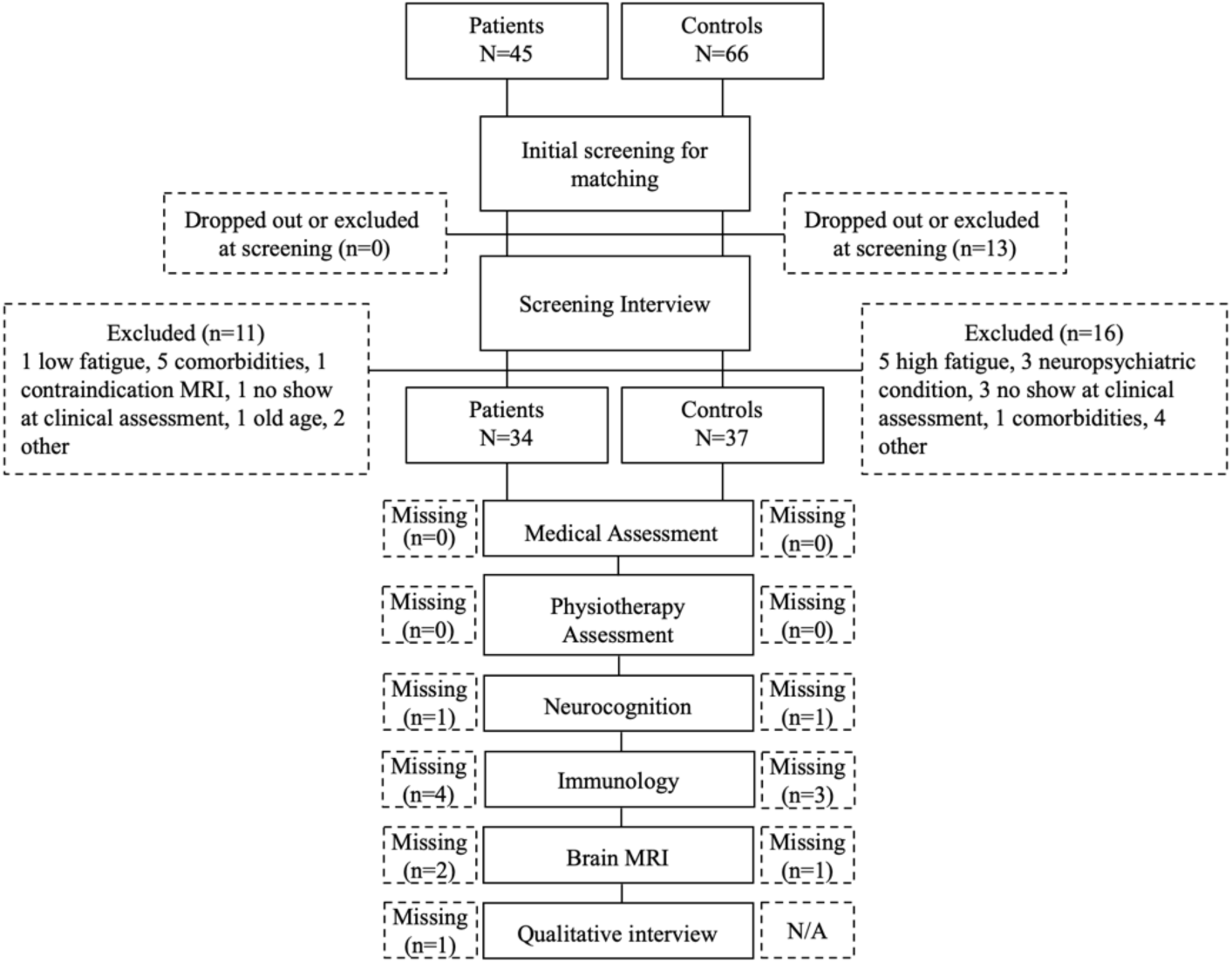
Flow chart of inclusion process.

### 2.3. Assessments

A summary of all assessments is provided below; for full methodological details, please refer to the study protocol^19^ and Supplementary Appendix.

#### 2.3.1. Medical and Physiotherapy Assessment

Each participant met with a physician who conducted a standard clinical examination, including a detailed medical history with a focus on COVID-19 and any persistent symptoms. Blood and urine samples were collected prior to clinical assessment. The PCC group also underwent functional and respiratory testing and completed corresponding self-reported questionnaires.

#### 2.3.2. Neuropsychological Assessment

A comprehensive neuropsychological test battery was administered to assess four cognitive domains. Attention was assessed using the Ruff 2 & 7 and CPT-III Detection. Working memory was assessed using a composite score derived from the forward and backward Digit Span subtests of the WAIS-IV. Processing speed was assessed using Coding (WAIS-IV), Trail Making Test A (TMT-A), and a composite score from Color Word Interference Test (CWIT) conditions 1 and 2, here referred to as CWIT processing speed. Executive function was assessed using Trail Making Test B (TMT-B), along with a composite score from CWIT conditions 3 and 4, here referred to as CWIT executive function. Additional measures included the Hospital Anxiety and Depression Scale (HADS) and the Insomnia Severity Index (ISI).

#### 2.3.3. MRI

Brain imaging was performed using a 3T Siemens Prisma scanner with a 64-channel head coil. Sequences included: Anatomical: T1-weighted and T2FLAIR images; functional: Resting-state fMRI (rs-fMRI); diffusion: multi-shell diffusion imaging; and relaxometry with Qmap sequence.

#### 2.3.4. Immunological Assessment

Plasma samples were analyzed using a Luminex™ multiplex assay to quantify selected cytokines, chemokines, and vascular markers relevant to post-COVID inflammation.

#### 2.3.5. Qualitative Phenomenological Analysis of Interviews

The qualitative phenomenological philosophy analysis centered on modal alterations, i.e. alterations in overarching ways of being, experiencing and engaging with others and the world, in research participants’ narrated lived experiences of fatigue in PCC. This analysis was performed on the phenomenological interview materials as a whole and specifically focused on experiences of cognitive difficulties.

### 2.4. Statistics and analyses

Measures were selected a priori based on theoretical and empirical considerations and hypothesized cross-domain relationships, while allowing for exploratory analyses across modalities, including immunological and neurocognitive variables.

Raw scores from each neuropsychological test were converted to age-adjusted scaled scores and T-scores in accordance with test manuals and normative, age-stratified reference data. To ensure comparability, scaled scores were transformed into T-scores (mean 50 SD 10). Higher scores indicate better neurocognitive performance. Group differences in neurocognitive performance and brain tissue volumes were examined using independent-samples *t-*tests. For Ruff 2 & 7 speed, TMT-A, TMT-B, and the CWIT composite scores, Levene’s Test indicated heteroscedasticity; therefore, adjusted degrees of freedom were applied.

Resting-state fMRI data were analyzed using MELODIC ^20^ to derive resting-state brain networks. Group analyses compared controls and PCC participants and examined associations between brain networks and neurocognitive test scores. Group comparisons were also performed for metrics computed from the diffusion weighted images, using tract based spatial statistics (TBSS) framework.^21^

Pearson correlation coefficients were calculated to assess associations among neurocognitive performance, immunological parameters, and MRI-based volumetric measures of white and gray matter, myelin, and brain parenchymal fraction (BPF).

Interaction analyses, between resting state fMRI, group and neurocognitive performance were performed using linear regression. Interaction analyses were also performed between diffusion MRI metrics, group, and immunology biomarkers.

### 2.5 Ethics

The Swedish Ethical Review Authority approved the study (Dnr 2022-02221-01/ 2022-03823-02) prior to recruitment. All participants provided informed written consent, after having been given written and oral information about the study.

## 3. Results

We first summarize group comparisons across MRI measures, neurocognitive performance, and immunological markers, together with the qualitative phenomenological analysis; detailed results are provided in the Supplementary Materials. We then report quantitative correlation analyses, beginning with pairwise associations among MRI, neurocognitive, and immunological measures followed by interaction analyses between MRI and neurocognitive performance, and between MRI and immunological markers. Finally, we relate neurocognitive performance to the qualitative phenomenological findings.

### 3.1. Participant characteristics

The PCC and the control group did not differ in key demographic variables, including age, sex, and years of education. Descriptive statistics for sex, age, educational level, and scores on fatigue, anxiety, and depression are presented in Table 1. Medical assessment, including scrutiny of medical records, confirmed eligibility criteria and revealed no clinically significant somatic, neurological, or routine laboratory abnormalities in either group.

**Table 1.** Sociodemographic factors.

|  | PCC (*n*=34) |  | Controls (*n*=37) |  | *t* | *df* | *p* |
| --- | --- | --- | --- | --- | --- | --- | --- |
|  | *m* | *sd* | *m* | *sd* |  |  |  |
| Age | 46·1 | 12·1 | 47·0 | 11·3 | -.328 | 69 | ·744 |
| Education (y) | 14·9 | 2·7 | 16·0 | 2·4 | -1·7869 | 69 | ·079 |
| HADS depression (*n*=31/*n*=36) | 6·58 | 3·49 | 2·25 | 2·64 | 5·768 | 65 | <·001 |
| HADS anxiety (*n*=31/*n*=36) | 5·71 | 3·02 | 4·17 | 2·67 | 2·219 | 65 | ·030 |
| MFI-20* (*n*=31/*n*=37) | 75·0 | 12·2 | 38·0 | 11·3 | - | - | - |
| FSS* (*n*=31/*n*=37) | 53·9 | 5·9 | 21·2 | 8·6 | - | - | - |
|  | *n* | % | *n* | % | - | - | - |
| Gender: men | 8 | 23·0 | 11 | 29·7 | - | - | - |
| Hospitalized | 10 | 29·4 | 1 | 2·7 | - | - | - |
| First language: Swedish | 30 | 88·2 | 36 | 97·3 | - | - | - |
| ISI ≥ 15 | 8 | 23·5 | 0 | 0 | - | - | - |

Note: HADS = Hospital Anxiety & Depression Scale, MFI-20 = Multidimensional Fatigue Inventory, FSS= Fatigue Severity Scale, ISI = Insomnia Severity Scale·
\*inclusion/ exclusion criteria are based on MFI-20, and FSS is highly correlated to MFI-20·

### 3.2. Group comparisons across modalities

#### 3.2.1. Functional and respiratory status

Participants with PCC and controls showed comparable performance on respiratory assessments; however, the PCC group demonstrated reduced physical capacity and reported higher symptom burden (see Supplementary Appendix).

#### 3.2.2. Neurocognitive results

Performance on all neurocognitive tests differed significantly between groups (Table 2), although mean scores in both groups were within the normal range (±1SD, see Supplementary Appendix).

**Table 2.** Correlations between neurocognitive performance, immunology and brain tissue volumes.

|  |  | RUFF2&7 | CPT-III<br>Detection | Coding | TMT-A | CWIT<br>Speed<br>Composite | TMT-B | Digit Span | CWIT<br>Executive<br>Composite | MMP1 | WFA2/AD<br>AMTS13 | MMP8 | WM | GM | MyC | V6 |
| --- | --- | --- | --- | --- | --- | --- | --- | --- | --- | --- | --- | --- | --- | --- | --- | --- |
| **Attention** | **RUFF2&7** | 1 | .366** | -.676** | -.518** | .362** | -.590** | .200 | -.537** | -.018 | -.061 | -.172 | .166 | -.006 | -.105 | -.028 |
|  | **CPT Detection** | .366** | 1 | .362** | .438** | .535** | .449** | .318* | .572** | -.141 | .206 | .000 | .245 | .142 | .152 | .033 |
| **Processing speed** | **Coding** | -.676** | .362** | 1 | -.630** | .459** | -.669** | -.317** | -.619** | -.051 | -.003 | -.082 | -.198 | -.126 | -.115 | -.270* |
|  | **TMT-A** | -.518** | .438** | -.630** | 1 | .530** | -.670** | .296* | -.544** | -.156 | -.021 | .131 | .313** | .159 | .268* | .101 |
|  | **CWIT Speed composite** | .362** | .535** | .459** | .530** | 1 | .516** | .393** | .699** | -.175 | .063 | .045 | .338** | .171 | .281* | .115 |
| **Executive function** | **TMT-B** | -.590** | .449** | -.669** | -.670** | .516** | 1 | -.394** | -.608** | -.076 | -.002 | -.019 | -.267* | -.108 | -.209 | -.206 |
| **Working memory** | **Digit Span** | .200 | .318* | -.317** | .296* | .393** | -.394** | 1 | -.475** | -.171 | -.118 | -.196 | -.294* | -.481** | .254* | -.285* |
| **Executive function** | **CWIT Executive Composite** | -.537** | .572** | -.619** | -.544** | .699** | -.608** | .475** | 1 | -.102 | -.076 | .021 | .317** | .242* | .253* | .138 |
| **Immunology** | **MMP1** | -.018 | -.141 | -.051 | -.156 | -.175 | -.076 | -.171 | -.102 | 1 | .364** | .251 | .034 | -.226 | .008 | -.101 |
| **Immunology** | **WFA2/ADAMTS13** | -.061 | .206 | -.003 | -.021 | .063 | -.002 | -.118 | -.076 | .364** | 1 | .040 | -.185 | -.259* | -.187 | -.189 |
| **Immunology** | **MMP8** | -.172 | .000 | -.082 | -.131 | .045 | -.019 | -.196 | .021 | .251 | .040 | 1 | -.017 | -.113 | -.007 | .040 |
| **MR** | **White matter** | .166 | .245 | -.198 | .313** | .338** | -.267* | .294* | .317** | .034 | -.185 | -.017 | 1 | .528** | .953** | -.287* |
| **MR** | **Grey matter** | -.006 | .142 | -.126 | .159 | .171 | .108 | .481** | .242* | -.226 | -.259* | -.113 | .528** | 1 | .568** | .383** |
| **MR** | **Myelin** | .105 | .152 | -.115 | .268* | .281* | .209 | .254* | .253* | .008 | -.187 | -.007 | .953** | .568** | 1 | .236 |
| **MR** | **Brain Parenchymal Fraction** | .028 | .033 | -.270* | -.101 | .115 | .206 | .285* | .138 | -.101 | -.189 | .040 | .287* | .383** | .236 | 1 |

Note: CPT Detect= Detectability measure from Conners Continuous Performance Test- 3<sup>rd</sup> Edition; TMT= Trail Making Test; CWIT Comp= Composite Score calculated as the mean scaled score of Inhibition and Inhibition/ Switching task from the Color-Word Interference Test; MMP= matrix metalloproteinase ;WFA2/ADAMTS13 = von Willebrand factor A2, ADAMTS: von Willebrand factor-cleaving protease.
\* = $p < .05$ , \*\* = $p < .001$

#### 3.2.3. MRI Assessments

Conventional structural MRI showed no between-group differences, with findings within the normal range. Relaxometry indicated lower white matter and myelin volumes in the PCC group compared with controls. Diffusion MRI revealed a focal area of reduced non-Gaussanity in the right superior/middle temporal gyrus in PCC (Supplementary Appendix Figure S1). The PCC group also showed higher functional connectivity across several brain networks (see Supplementary Appendix Figures S2 – S4).

#### 3.2.4. Cytokine Profile

In a multiplex immunoassay of 17 inflammatory mediators, three mediators (MMP1, MMP8 and the VWF A2/ADAMTS13 ratio) were found to be significantly induced before, but not after, correcting for multiple comparisons (Supplemental Appendix Table 3).

#### 3.2.5. Summary of the Qualitative Phenomenological Analysis

Two modal alterations were identified:

i. i. an alteration in which participants in the PCC-group experienced difficulties to be present in the “here and now”, concentrate and “hold” a thought and/or follow a “thread” in thinking and conversations, and correspondingly experienced themselves as “zooming out” and, across all these experienced difficulties, an altered sense of self (referred to as *Difficulties to concentrate or follow a “thread”*, see Supplementary Appendix), and
ii. ii. an alteration in some of these participants’ experienced abilities to execute decisions and actions, which correspondingly involved an altered sense of selves, as no longer “themselves”, as “slow” or incapable (referred to as *Difficulties to execute decision and actions)*.

As some examples, (i) was expressed in narrations of how following a topic of conversation “pours away from me” or of referring to oneself as becoming “blank”, while (ii) was reflected in narrated experiences not only of having difficulties to execute decisions and actions, generally, but also more specifically of no longer being able to execute decisions and actions tacitly and pre-reflectively. Instead, such execution now required explicit attention to how to act. While 26 of the interviewees narrated experiences corresponding to (i), half of them also shared experiences corresponding to (ii). None of them told only about the second kind of experience. Phenomenologically, cognitive and affective dimensions are intertwined at the experiential level, and the analysis brought out a basic “bodily doubt”, which involves a thorough disruption of one’s sense of certainty, trust, and confidence in one’s own body and bodily capacities.^22(p.184^)

### 3.3. Correlations between neurocognitive performance, immunology and brain tissue volumes

Significant correlations were observed between immunological markers and fatigue measures in the full sample. MFI-20 scores were positively associated with MMP1, (r = ·312, p = ·013), as were FSS scores and MMP1, (r = ·269, p = ·034). When analyses were restricted to the PCC group (n = 28), none of these associations remained statistically significant. In the control group (n = 34), significant positive correlations were observed between anxiety and MMP1, (r = ·423, p = ·014), as well as between MFI-20 and MMP1, (r = ·464, p = ·006). No significant associations were found between MRI variables and fatigue, anxiety, or depression.

Analyses of associations between neurocognitive performance and immunological markers revealed no significant correlations. In contrast, neurocognitive performance was significantly associated with MRI-derived brain metrics. White matter, grey matter, myelin content and brain parenchymal fraction (BPF) were positively correlated with processing speed (Coding and TMT-A), executive functioning (TMT-B and CWIT executive composite scores), and working memory (Digit Span). Larger brain tissue volumes were associated with better performance. These associations were attenuated in the PCC group, although several remained statistically significant (Table 2).

A significant negative correlation was observed between immunological markers (the VWF A2/ADAMTS13 ratio) and gray matter volume (r = –·259, p = ·040, n = 63). No other correlations between immunological and MRI metrics were statistically significant. In the PCC group, this association did not reach significance (r = –·352, p = ·052, n = 31). No significant associations were identified in the control group.

### 3.4. Pairwise cross-modal interaction analyses

#### Interactions between neurocognitive performance and rs-fMRI connectivity

Interaction analyses between brain network connectivity, group status and neurocognitive test performance (Coding, Ruff 2&7, CWIT executive composite score, CWIT processing speed composite score) identified several significant interactions (p < ·05, FWE corrected within each brain network).

Within the dorsal attention network (DAN) significant interactions were observed in the prefrontal gyrus and intraparietal sulcus regions for CWIT executive composite scores (Figure 2). In the salience network, a significant interaction with coding was observed in the insula (Figure 3). In the left executive network, a significant interaction with CWIT executive composite score was identified in the dorsolateral prefrontal cortex (Figure 4).

**Figure 2.**
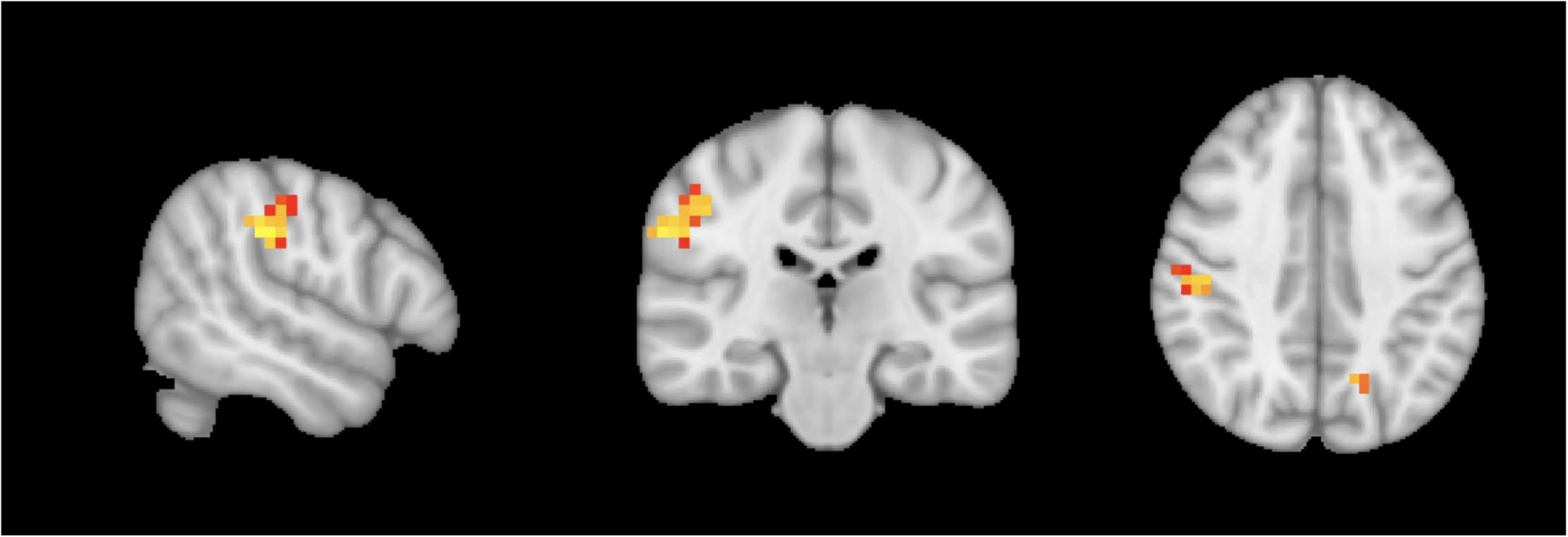
Significant interaction between dorsal attention network (DAN) connectivity, group and CWIT executive composite scores. Separate group analyses suggest that controls have negative correlation between DAN and CWIT in the intraparietal sulcus (see Supplementary Figures S5, S6), which is not observed in the PCC group. The separate group analyses also suggest that the PCC group has a positive correlation between DAN and CWIT in the intraparietal sulcus (see Supplementary Figures S5, S6), which is not observed in controls. Results are based on N = 65, thresholded at p = ·05, FWE-corrected using TFCE. The lowest corrected p-value was ·0052. Red indicates that the p-value is close to p = ·05, while yellow indicates a lower p-value.

**Figure 3.**
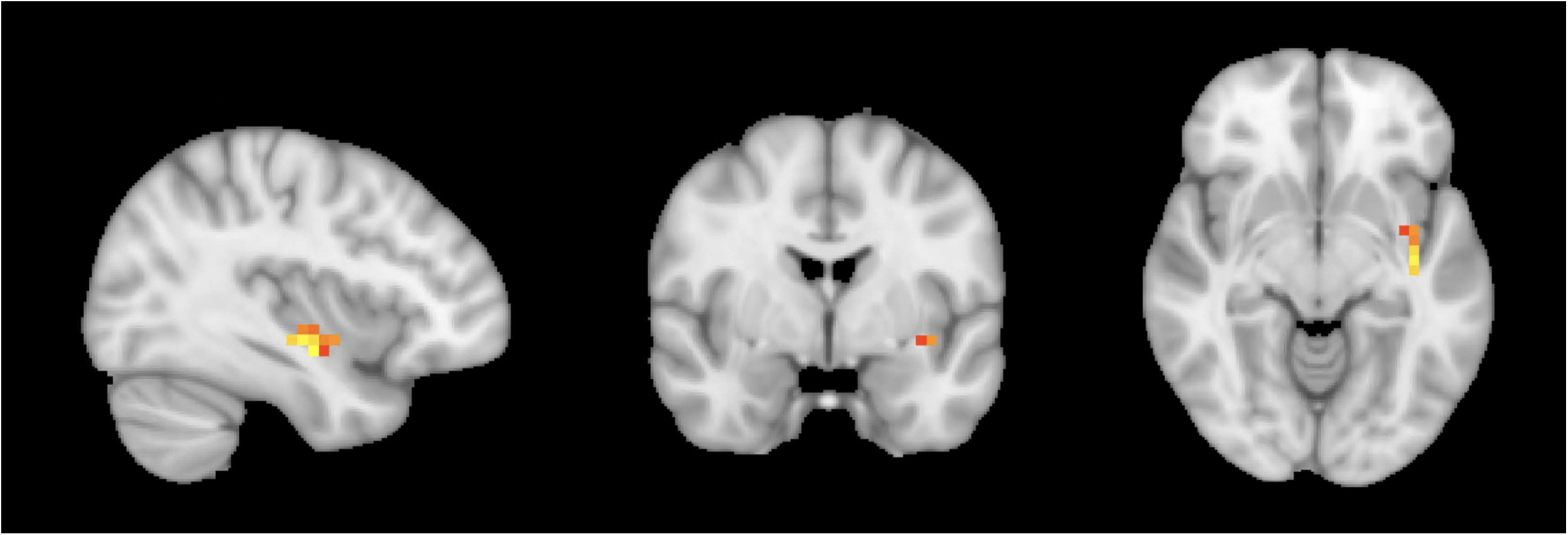
Significant interaction between salience-network connectivity, group and performance on the Coding test. Separate group analyses suggest that the PCC group has a negative correlation between salience-network connectivity and Coding (see Supplementary Figure S7), which is not observed in controls. Results are based on N = 65, thresholded at p = ·05, FWE-corrected through TFCE. The lowest corrected p-value was ·024. Red indicates that the p-value is close to p = ·05, while yellow indicates a lower p-value.

**Figure 4.**
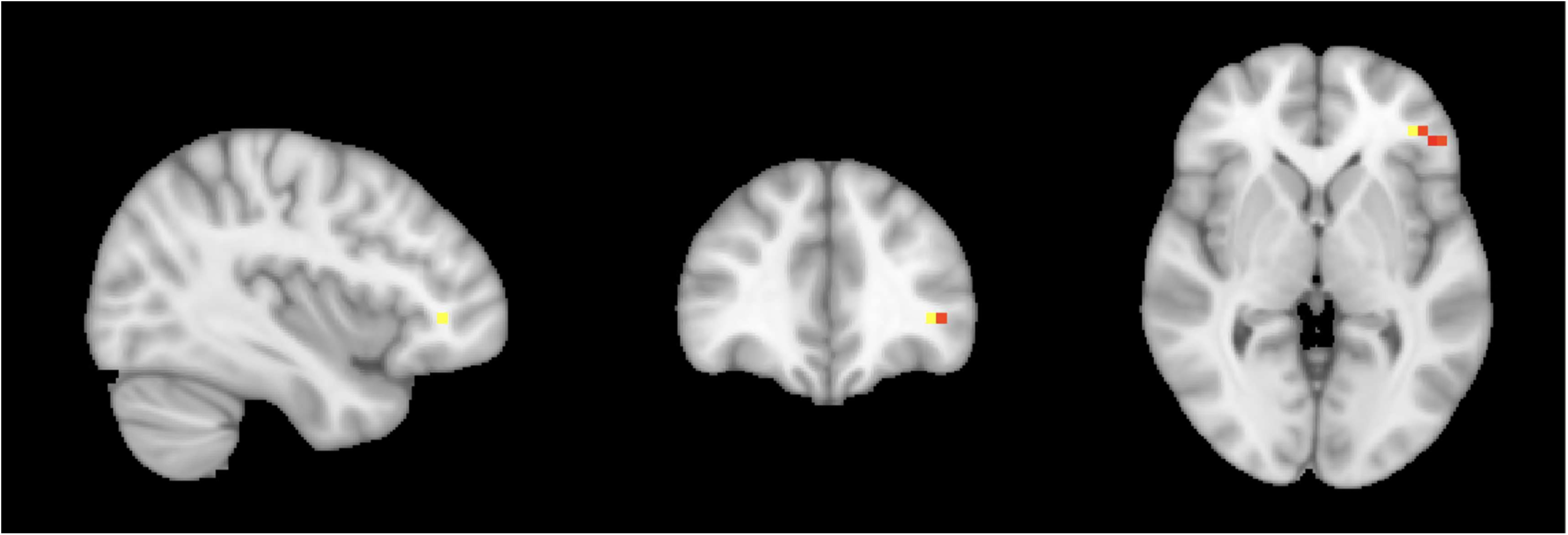
Significant interaction between left central executive network connectivity, group and CWIT executive composite scores. Separate group analyses suggest that controls have a negative correlation between left frontoparietal network and CWIT executive performance (see Supplementary Figure S8), which is not observed in the PCC group. Results are based on N = 65, thresholded at p = ·05, FWE-corrected through TFCE. The lowest corrected p-value was ·0282. Red indicates that the p-value is close to p = ·05, while yellow indicates a lower p-value.

Interactions in the DAN and the salience network remained significant in two-sided analyses. Total sample size for the analyses was N = 65 due to missing neurocognitive data.

##### 3.4.1. Interactions between immunological markers and diffusion MRI measures

Interaction analyses between diffusion MRI metrics (fractional anisotropy (FA), mean diffusivity (MD), mean squared displacement (MSD), non-Gaussanity (NG), return to origin probability (RTOP), and Q-space inverse variance (QIV)), group status, and immunology markers (MMP1, MMP8 and the VWF A2/ADAMTS13 ratio) identified several significant interactions (p < 0·05, FWE corrected).

For RTOP, significant interactions with MMP1 were detected across widespread white-matter regions (Figure 5). For MD, a significant interaction with VWF A2/ADAMTS13 ratio was found in the precuneus (Figure 6), although this effect did not remain significant in two-sided analyses. For the remaining diffusion metrics, the lowest interaction p-values were: FA with MMP1, p =0·0606; MSD with the VWF A2/ADAMTS13 ratio, p = 0·0502; and QIV with the VWF A2/ADAMTS13 ratio, p = 0·0542. Analyses were based on N = 53 due to missing immunology data.

**Figure 5.**
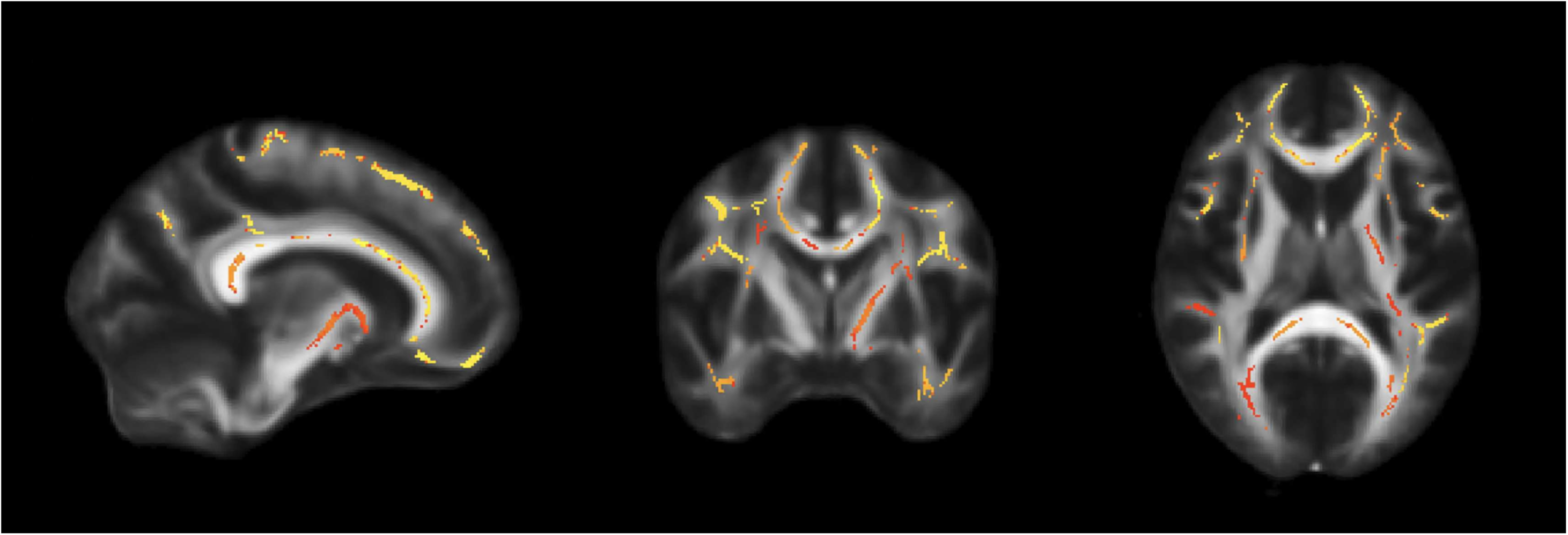
Significant interaction between the diffusion metric RTOP (return to origin probability), group, and MMP1. This interaction may reflect a more widespread positive correlation between RTOP and MMP1 in the controls, while the PCC group has a widespread negative correlation (see Supplementary Figures S9, S10). Results are based on N = 53, thresholded at p = ·05, FWE-corrected through TFCE. The lowest corrected p-value was ·0214. Red indicates that the p-value is close to p = ·05, while yellow indicates a lower p-value.

**Figure 6.**
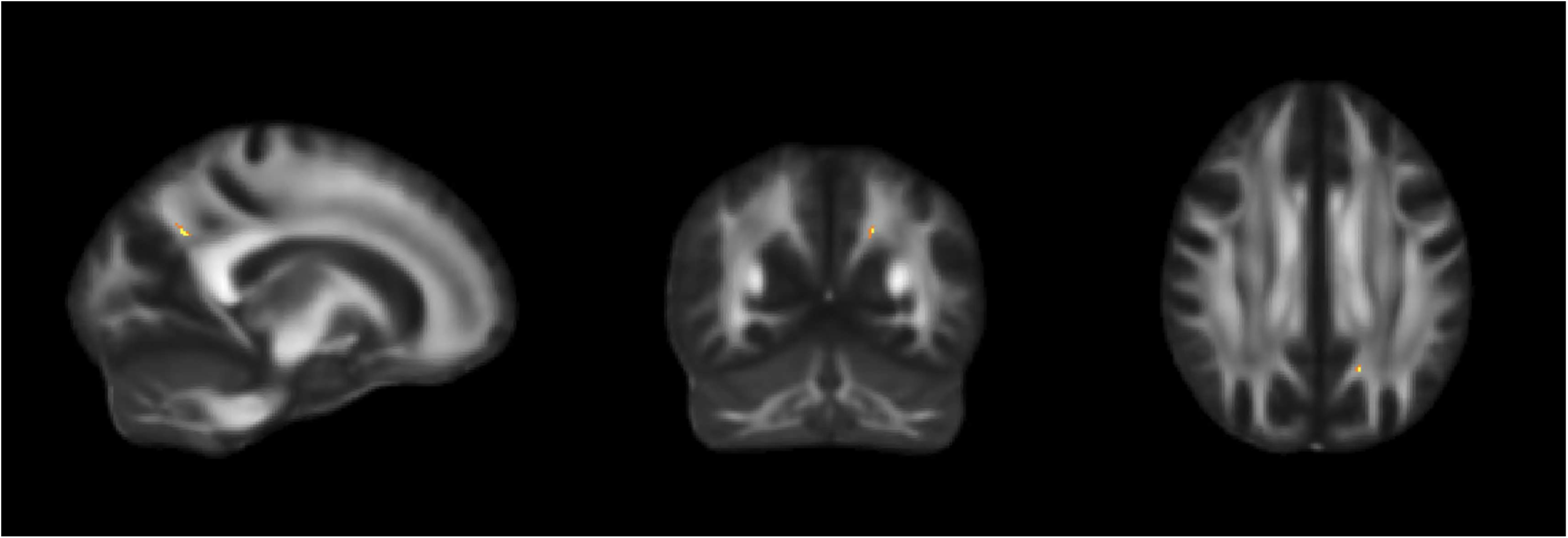
Significant interaction between mean diffusivity (MD), group and WFA2. Separate group analyses indicate that the PCC group has a widespread negative correlation between MD and WFA2 (see Supplementary Figure S11), which is not observed in controls. Results are based on N = 53, thresholded at p = ·05, FWE-corrected through TFCE. The lowest corrected p-value was ·0496.

##### 3.4.2. Phenomenology in relation to neurocognitive performance

While not designed for direct comparison at the individual level, the identified modal alterations can be described in relation to the neurocognitive domains assessed. Difficulties in ‘following a thread’ or ‘holding on to’ a line of thought may be interpreted as resonating with functions such as sustaining attention, working memory and processing speed, capacities that the Coding task, Ruff 2 & 7, and the CWIT place demands on. Similarly, reported difficulties in executing decisions and actions, and the shift from tacit to reflective action corresponds to aspects of executive control and reduced automaticity.

## 4. Discussion

In this interdisciplinary study of PCC, we combined brain MRI, neurocognitive, and immunological data, and qualitative phenomenological analysis of interviews to examine the biological, cognitive, and experiential dimensions of persistent cognitive symptoms in individuals with PCC and fatigue compared with matched controls. Findings across modalities suggest a pattern of subtle white matter and network-level vulnerability, rather than cortical volume loss. These alterations appear sufficient to reduce cognitive efficiency in a subtle manner, despite neurocognitive performance remaining within the normal range, consistent with findings of difficulties in sustaining attention and executing actions identified in the qualitative phenomenological analysis.

This pattern aligns with previous studies linking persistent cognitive symptoms in PCC to alterations in brain networks and white-matter organization rather than to cortical atrophy.^14,23^ Neurocognitive performance showed consistent associations with both structural MRI metrics and functional network connectivity. Processing speed, executive function, and working memory were associated with brain tissue volumes, with several associations remaining significant when analyses were restricted to the PCC group. Similar associations between grey matter alterations and cognitive dysfunction have been reported in previous post-COVID studies.^11,15^

At the network level, increased activity in the salience and dorsal attention networks were associated with lower performance on tasks of processing speed and attention and interaction effects, indicating differing patterns of association across groups. These associations suggest differences in network engagement in individuals with greater cognitive difficulty, potentially reflecting reduced neural efficiency or compensatory activation. This interpretation is supported by previous PCC studies reporting altered or inefficient engagement of attention-related and default-mode networks in individuals with cognitive complaints and fatigue.^12,14,23^ Interaction effects within the dorsal attention and executive network indicate that task-control systems are engaged more strongly in PCC despite lower executive performance. This dissociation between network engagement and behavioral output suggests altered regulation of goal-directed control processes rather than loss of core executive capacity also discussed by Schumann and colleagues.^24^

Our findings suggest that the functional network alterations observed in PCC may be related to compensatory network reorganizations in response to underlying microstructural white matter alterations and immunological processes. This interpretation is supported by the interaction between immunological biomarkers (MMP1) and advanced microstructural diffusion MRI metrics (RTOP), which indicate opposing associations in patients and controls (Supplementary Appendix Figures S9 and S10). The absence of pronounced group differences in diffusion and immunology measures suggest heterogeneity rather than robust differences in individual modalities. As in previous studies, we performed group-wise comparisons of standard diffusion metrics (e.g. FA, MD),^25,26^ but we identified limited group differences. In contrast, interaction-based analyses combining advanced diffusion metrics and immunological markers identified cross-modal associations not evident at the group level. While cytokines have been linked to white matter differences in several neurological diseases,^27,28^ such findings remain limited in PCC.

While the quantitative findings showed largely preserved performance on standard cognitive tests in individuals with PCC, the qualitative phenomenological analysis of interviews highlighted experienced difficulties to be present in the “here and now”, to concentrate, and execute decisions and actions, which correspondingly entailed an altered sense of self. This analysis underlines existential dimensions, such as distinct felt losses: participants could no longer engage with others and the world as before, combined with a sense of oneself as no longer “oneself.” Such loss was also combined with a basic bodily doubt,^22^ not only in one’s own abilities, but also whether one might regain previously taken-for-granted ways of being, experiencing and engaging with others and the world. Whitin the present study, the basic bodily doubt appeared as centered on the self’s own cognitive faculties – i.e., on faculties that are deeply involved in the everyday tacit assessing of one’s capacities to engage with others and the world (see Supplementary Appendix).

The phenomenological analysis identified difficulties in “following a thread”, sustaining attention, and executing actions with fluency, including a shift from tacit, pre-reflective action to a more reflective and effortful mode. However, these findings cannot readily be reduced to discrete cognitive functions; rather, they point to broader alterations in how the self relates to its lived world. When read alongside the neurocognitive findings, these distinct emphases suggest how subtle neural changes may be experienced in everyday cognitive functioning in PCC, particularly under conditions of fatigue. The dissociation between preserved cognitive test performance and subjective cognitive difficulties, aligns with prior work distinguishing cognitive capacity from the accessibility and sustainability of cognitive performance, particularly under conditions of fatigue.^4^ This dissociation suggests that standard measures may underestimate meaningful shifts in cognitive functioning.

Taken together, the phenomenological and neurocognitive findings capture complementary aspects of cognitive change. The former reflects lived experiences of losing the thread, becoming unanchored, and losing the tacit ease of action, while the latter indicates subtle but consistent reductions in processing speed, sustained attention, and executive efficiency despite performance remaining within normal ranges. This convergence of pronounced experiential changes and subtle objective findings underscores the value of integrating these approaches to more fully characterize cognitive alterations.

Participants with higher levels of systemic immune activation reported greater fatigue, consistent with evidence that inflammatory processes may contribute to perceived fatigue.^3,29^ In the control group, additional associations were observed between anxiety and fatigue measures and one immunological marker. Although MMP1 and MMP8 levels, and the VWF A2/ADAMTS13 ratio largely overlapped between PCC and control group, they may still hold biological relevance for certain individuals. This is supported by our finding of a negative correlation between MMP1 and RTOP in individuals with PCC, as MMPs are known to influence white-matter microstructure.^30^

Fatigue presents with similar symptom profiles across multiple conditions, including multiple sclerosis, rheumatoid diseases, mild traumatic brain injury, and exhaustion disorders.^24^ Our findings are broadly in line with this literature, as we found no evidence of a distinct pattern specific to post-COVID fatigue, instead pointing toward inefficient or compensatory network engagement as a potential transdiagnostic mechanism, and highlighting the difficulty of distinguishing fatigue based solely on cognitive or neuroimaging measures.

### 4.1. Strengths and limitations

One strength of this study is the inclusion of participants with a clinically established diagnosis of PCC, rather than recruitment based solely on symptom self-report. An additional strength is the inclusion of a demographically matched control group with a confirmed history of COVID-19, allowing comparisons between groups matched for education level, age, and sex, all of which are known to influence cognitive performance. A major strength is the integration of multiple research fields, combining neurocognitive testing, neuroimaging, immunological measures, and qualitative phenomenological analysis. This multimodal approach allowed cognitive difficulties to be examined from complementary perspectives within a single cohort.

Several limitations should nevertheless be acknowledged. Due to the cross-sectional design, it is not possible to determine whether observed cognitive differences reflect changes emerging after COVID-19 infection or pre-existing vulnerability. Although statistically significant group differences were identified across several neurocognitive measures, relatively few individuals with PCC met conventional criteria for objective cognitive impairment. This underscores the limited sensitivity of standardized assessments for detecting subtle or fatigue-related cognitive difficulties, particularly when performance remains within normative reference ranges.

In both PCC and control groups, performance on neurocognitive tests and level of immunological markers largely fell within reference ranges. Group differences primarily emerged in MRI findings, the interaction analyses, and in the cross-modal analyses. As phenomenological interviews were conducted only in the PCC group, we cannot determine how control participants would have narrated experiences of fatigue and cognitive functioning. This limits conclusion regarding the specificity of the narrated experiences from a biomedical perspective.

Despite exclusion criteria intended to minimize the impact of conditions known to affect cognition or fatigue, the PCC group exhibited a higher burden of comorbid diagnoses compared with controls. Such premorbid factors may have contributed to variability in cognitive outcomes and should be considered when interpreting group differences.

In addition, while the sample size was very large for a qualitative phenomenological analysis, it may have limited statistical power to detect small but clinically meaningful effects in the quantitative analyses. Furthermore, given the number of statistical tests performed, several of the significant interaction effects may not remain significant after correction for multiple comparisons. Finally, SARS-CoV-2 infection status was based on self-reported positive test results and was not independently verified through medical records.

## 5. Conclusion

This study combines neuroimaging, neurocognitive assessment, immunological measures, and qualitative phenomenological analysis within a single cohort, enabling cross-modal analysis of persistent cognitive symptoms in PCC. This holistic approach allows subjective, lived experience and objective system-level measures to be interpreted in relation to one another.

Taken together, and within the limitations of a small, fatigue-enriched sample, our results point toward a pattern in which cognitive difficulties in PCC may be better understood in terms of reduced network efficiency and cognitive endurance rather than loss of cognitive capacity.

The convergence of pronounced experiential alterations and subtle objective findings suggests that white-matter and large-scale network organization changes may relate to altered modes of experiencing and acting. While this interpretation remains tentative and hypothesis-generating, it aligns with emerging views of PCC as a condition characterized by systems-level vulnerability that becomes evident under sustained cognitive and physiological load. Meaningful cognitive alterations in PCC may therefore not be adequately captured by standard measures alone, but emerge most clearly when experiential and clinical objective perspectives are considered together within an integrative framework.

## Contributor statement

Conceptualization: all authors; data curation: UBT, AA, AE, KZ, SMJ, RL, DE, AD, IB; formal analysis: UBT, AE, KZ, SMJ, FL, RL, DE, AD, IB; funding acquisition: KZ, RL, investigation: UBT, AE, KZ, SMJ, FL, AA, RL, DE, AD, IB; methodology: all authors; project administration: KZ, AA, RL, IB; resources: AE, KZ, RL, DE, IB; software: UBT, AE, DE, IB; validation: UBT, AE, KZ, SMJ, RL, DE, AD, IB; visualization: UBT, AE, AA, AD, IB; writing – original draft: UBT, AE, KZ, IB; and writing – review & editing: all authors.

## Funding

Swedish Research Council (Vetenskapsrådet) (Dnr 2021-01245). IB holds a grant as an associated clinical fellow with the Wallenberg Center for Molecular Medicine.

## Role of the funding source

The funding body had no role in study design, data collection, data analysis, data interpretation, or writing of the manuscript.

## Declaration of interest

Authors report no conflict of interest.

## Data sharing

The data presented in this study are not publicly available due to the ongoing nature of the project, privacy considerations, and the Swedish ethical regulations and approvals for the project. The study protocol is available.^19^

## Acknowledgements

We thank all the research participants for participating in the study. We also would like to thank Johanna Wibault at Rörelse och Hälsa, Linköping University Hospital, and Mirjam Schimanke at Norrköping Hospital and their clinical teams for their assistance with the recruitment of research participants; the team at CMIV for support with the MRI exams, Silvia Castany for performing the Luminex-based analysis; the clinical team at the Clinical Department of Pain and Rehabilitation Medicine at Linköping University Hospital for providing input in the design of the clinical assessment; especially the physiotherapists for conducting the physical examination of patients and controls and the psychologists for assisting UBT and AA with performing the neuropsychological assessments; as well as Therese Dossel Holm for her work with administration of clinical processes.

## Notes

### Competing Interest Statement

The authors have declared no competing interest.

