## Supplemental material for "Understanding cognitive function in post-COVID-19 condition with fatigue: an interdisciplinary multimodal investigation"

### Supplement

**1. Methods**

**1.1 Medical assessment**

**1.2 Physiotherapy assessment**

**1.3 Neuropsychological assessment**

**1.4 Brain MRI**

**1.5 Immunological analysis**

**1.6 Qualitative Phenomenological analysis**

**2. Results**

**2.1 Medical assessment**

**2.2 Physiotherapy assessment**

**2.3 Neuropsychological assessment**

**2.4 Brain MRI**

**2.5 Immunological results**

**2.6 Qualitative Phenomenological Analysis**

**2.7 Correlation analysis**

#### 1. Methods

##### 1.1 Medical assessment

The PCC group as well as controls underwent a thorough clinical examination by a physician specialized in rehabilitation medicine. The examination comprised scrutiny of previous medical records, taking and documenting the medical history regarding previous and current health issues and comorbidities, initial and persisting symptoms related to COVID-19 (including severity of initial infection), somatic and neurologic physical examination, and assessment of routine laboratory test results. For details, please see Levi et al [1]. The rationale for medical assessment was primarily to confirm previous COVID-19 and the temporal relationship of current PCC-related symptoms to this infectious event. Further, to exclude any comorbid conditions of such type and/or severity to likely influence the current symptoms.

##### 1.2 Physiotherapy assessment

Pulmonary function was assessed via the modified Medical Research Council (mMRC) scale; Spirometry (VC, FEV1, PEF) using Micro-Lab ultrasonic spirometer; Maximal inspiratory and expiratory pressures (MIP/MEP) via MicroRPM, with reference values from Evans & Whitelaw; Screening for dysfunctional breathing (Nijmegen questionnaire); Physical activity level via the Grimby and Frändin 6-level scale; aerobic capacity and endurance via the 6-minute Walk Test (6MWT); functional fitness via the 1-minute sit-to-stand test (1mSTS).

##### 1.3 Neuropsychological assessment

Neurocognitive functioning was evaluated using a 2-hour comprehensive face-to-face neuropsychological assessment. The testbattery included multiple measures for each cognitive domain: attention, working memory, processing speed, and executive function.

Attention was assessed with the Ruff 2&7 [2] and the computerized Conners Continuous Performance Test, 3rd edition (CPT-III) [3]. Working memory was evaluated using forward and backward digit span composite scores from the Wechsler Adult Intelligence Scale, 4$^{th}$ edition (WAIS-IV)[4], as well as the Rey Auditory Verbal Learning Test[5].

Processing speed was measured using The Coding subtest from WAIS-IV,Trail Making Test A [6] and a processing speed composite score from the Color-Word Interference Test (D-KEFS) [7], calculated as the mean of conditions 1 and 2.

Executive function was assessed using the Trail Making Test B [6] and an executive composite score from the Color-Word Interference Test (D-KEFS) [7], calculated as the mean of conditions 1 and 2.

Cognitive domains were constructed based on standardized T-scores derived from the raw neuropsychological test scores. Depression and anxiety were assessed with the Hospital Anxiety and Depression Scale (HADS)[7] as a brief measure for depression and anxiety levels. Sleep disturbances were assessed with The Insomnia Severity Index (ISI)[8].

##### 1.4 Brain MRI

###### Image acquisition

The brains of the participants were examined in a 3T MR scanner (Prisma, Siemens, Erlangen, Germany) using a 64-channel head coil. In addition to conventional MR images of the brain, experimental sequences assessing tissue relaxation, diffusion, and functional connectivity were acquired. The MR protocol is reported in [1].

###### Neuroradiological assessment

Conventional images were analyzed by an experienced neuroradiologist and the white matter assessed with Fazeka's scoring [9].

###### Relaxometry

Relaxometry data were computed with SyMRI 15.0 Beta 1 (SyntheticMR AB, Linköping, Sweden) for tissue and myelin volumes and analyzed with SPSS: independent samples t-test for groupwise comparison.

###### Diffusion MRI

The DICOM files were converted to nifti using dcm2niix. The dMRI data were analyzed with FSL 6.0.2 [10], dipy 1.9.0 [11], *MRtrix3* [12] and Matlab 2019b. Data were corrected for head motion, susceptibility distortions and eddy currents using the functions eddy [13] and topup in FSL. For conventional dMRI analysis, volumes with a b-value of less than 1500 were used to fit a diffusion tensor model in each voxel, using a Matlab implementation (https://github.com/DenebBoito/qtiplus), which features a sequence of linear and non-linear fits with a positivity constraint on the estimated diffusion tensor [14–16]. After fitting the diffusion tensor, the scalar metrics FA (fractional anisotropy), MD (mean diffusivity), RD (radial diffusivity), and AD (axial diffusivity) were calculated from the diffusion tensor in each voxel.

To elucidate the information made available by the heavily diffusion-weighted acquisitions, the more advanced mean apparent propagator (MAP-MRI) technique [18] was employed, using the implementation in dipy [12] and functions in nibabel. The fitting was done for a radial order of 6 using the MOSEK optimizer, and global positivity constraints were applied [19]. In the computations for the scale matrix, a b-value threshold of 1500s/mm$^2$ was used. After the fitting, the scalar metrics MSD (mean square displacement), QIV (q-space inverse variance), NG (non-Gaussianity), and RTOP (return to origin probability) were computed. The transformations obtained from the TBSS analysis on FA were applied to these MAP-MRI scalar metrics to be able to perform group analyses within the white matter skeleton. Group analyses were performed using tract based spatial statistics (TBSS) [20] and randomise (i.e. permutation testing) [21].

Interaction analyses included covariates for MMP1, WFA2 and MMP8, a group membership covariate, interaction covariates for MMP1, WFA2 and MMP8 and an intercept. Correlation analyses per group included covariates for MMP1, WFA2, MMP8 and an intercept. All group analyses included covariates for age and sex. Inference was done using threshold free cluster enhancement [21] adapted for TBSS.

**Resting-state functional MRI (rs-fMRI)**

The DICOM files of the rs-fMRI sequence were converted to nifti using dcm2niix. The rs-fMRI data were analyzed with group ICA in FSL 6.0.2 [10] using MELODIC 3.15 [22]. The five first volumes (out of 800) were deleted for each subject, and a high pass filter with a cutoff of 100 seconds was used. Head motion correction was performed for each subject using MCFLIRT [23]. No subject was removed due to excessive head motion. Five mm FWHM spatial smoothing was applied to each volume. Slice timing correction was applied to correct time differences between slices, using a custom slice timing file. Each anatomical T1w volume was segmented using BET [24], to obtain brain-extracted volumes. All fMRI data were registered to MNI152 space using the anatomical T1w volumes and a combination of boundary-based registration [25] and linear registration with 12 degrees of freedom [23]. The group ICA was performed using multi-session temporal concatenation, with 40 output ICA components.

Group analyses were performed using dual regression [26] and randomise (i.e. permutation testing) [20]. To only perform the group analyses within each resting state network, instead of full brain analysis, the group ICA map for each resting state network was thresholded at z = 2.5 and z = -2.5 to obtain a binary network mask for each network.

Interaction analyses included covariates for coding, RUFF, CWIT executive composite score, CWIT processing speed composite score, a group membership covariate, interaction covariates for coding, RUFF, CWIT executive composite score, CWIT processing speed composite score, a covariate for sex (but not for age due to age adjusted psychometric scores), and an intercept. Correlation analyses per group included covariates for coding, RUFF, CWIT execution composite score, CWIT processing speed composite score, sex and an intercept. Group comparisons included covariates for both age and sex. Inference was done using threshold free cluster enhancement [21].

**1.5 Immunological assessment**

Following venous blood sampling, BD Vacutainer EDTA tubes were centrifuged at 2500g for 10 minutes. Plasma was transferred to a new tube and kept refrigerated during transport to the storage facility. There, samples were aliquoted and stored at -80°C. Plasma samples were thawed, and samples were transferred to 96 well plates for further analysis using a customized Luminex™ multiplex assay according to the instructions of the manufacturer. Briefly, 50 µL of diluted sample and 50 µL of magnetic, premixed, microparticle cocktail with antibodies specific for each mediator were mixed in each well and incubated in an orbital shaker. Samples were washed using a magnetic bead separator (HydroFlex™) and incubated with a premixed biotin-antibody cocktail with antibodies specific for each cytokine/chemokine. Antibody/analyte complexes were stained with Streptavidin-PE, and plates were read using a Luminex FLEXMAP 3D w/xPONENT 4.3 instrument. A standard curve was generated for each analyte to convert median fluorescence intensity to concentrations (typically pg/ml). Molecular targets were selected based on previous studies [27–36] and included ADAMTS13, APP, CCL4, CCL11, CCL19, CXCL5, Galectin-1, IFN-beta, IL-1 beta, IL-1 RII, IL-6, IL-7, IL-8, MMP-1, MMP-8, P-Selectin, TNF-alpha, and vWF-A2.

**1.6 Qualitative phenomenological analysis**

Analyses that integrate qualitative research and phenomenological philosophy are gradually becoming more common, especially in the field of phenomenology of medicine, illness, and health research (see [37,38]). Such analyses are performed with the aim of offering a deepened understanding of structures of experience such as embodiment or temporality and modal alterations, i.e., alterations in overarching, encompassing ways of being, experiencing and engaging with others and the world, in lived experiences of, for example, illnesses or bodily or functional variations.

The qualitative phenomenological analysis within this project was performed based on semi-structured phenomenological interviews with participants with PCC and fatigue. Each interview lasted between 60-70 minutes and was recorded and transcribed verbatim. Themes in the interview guide included: how the participant experienced fatigue; variations/dimensions of fatigue; experiences of oneself and one's relations with others when experiencing fatigue; experiences of how fatigue affects everyday life, and experiences of fatigue during the research process. The interview guide was phenomenologically "front-loaded" [37,39], i.e., basic phenomenological foci (such as embodiment) were included in it.

Further, as part of the work with the present interdisciplinary analysis, the research team as a whole (i.e. all co-authors) discussed the foci and analytic demarcations as part of the analysis process and the writing of the present article. This informed the choice of what to include and present in the qualitative phenomenological philosophy analysis presentation, in section 2.6, Supplement. The two themes presented in this article were main themes that centered on cognitive issues, and they were derived through the analysis of the interviews as a whole. Other themes have been presented elsewhere (see [40]).

The phenomenological interviews were performed by SMJ. The qualitative phenomenological analysis comprised an initial coding and clustering of codes within the material into subthemes, performed individually by KZ and SMJ. KZ and SMJ then compared and discussed their coding and the sub-themes, thereby ensuring a robust analysis (theme (i) subthemes: experiences of difficulties to hold a thought and or follow a thread in thinking and conversations; experiences of difficulties in processing stimuli; experiences of difficulties to be present in the present; experiences of being "slow" in one's thinking; sense of self as zooming out; altered sense of self as part of all these experiences. Theme (ii) subthemes: difficulties executing plans; difficulties performing actions; difficulties with initiatives; the need to think about how to perform previously taken-for-granted embodied skills; difficulties to take in stimuli and make quick decisions; altered sense of self as part of all these experiences).

In the next step, the coding and larger sections of the interviews were translated into English and discussed with FL. KZ, SMJ and FL clustered sub-themes with attention to overall modal alterations, and this step was followed by engagement with relevant phenomenological literature, and, in turn, a movement back and forth between the analysis of the interviews and phenomenological conceptual discussions and literature.

The number of phenomenological interviews, n=33, is large for these kinds of interviews, and data saturation was reached. As seen, 26 of the 33 interviewees narrated experiences corresponding to what we identify as the first theme (i) *Difficulties to concentrate or follow a "thread"*, see 2.6., Supplement), and 13 of these 26 also narrated experiences involving difficulties to execute decisions and actions, and an altered sense of themselves, i.e. what we identify as the second theme of *Difficulties to execute decision and actions*.

#### 2. Results

##### 2.1 Medical assessment

No participant included in the study presented any clinically remarkable findings as regards clinical somatic and neurologic examination; neither any pre- nor comorbid conditions likely to influence current symptomatology. Routine blood and urine tests were within normal limits, with occasional but clinically non-significant exceptions. However, the medical history obtained by medical assessment did confirm PCC-related symptoms, which received a more detailed scrutiny in the physiotherapeutic, neuropsychological, and phenomenological analysis, respectively. It may then be concluded that the medical assessment did not disclose any objective pathological findings but did confirm the inclusion and exclusion criteria of the study.

##### 2.2 Physiotherapy assessment

The PCC group demonstrated reduced physical capacity compared with controls, and a higher frequency of values below the lower limit of normal (LLN) on the 1mSTS (9 vs 1; $p = 0.005$). As compared to controls, the PCC group also had lower percent predicted values for both 6MWT (91% vs 100%; $p = 0.017$) and 1mSTS repetitions (89% vs 100%; $p = 0.014$).

Respiratory function was largely comparable between groups with respect to the proportion below LLN and percent predicted values; however, the PCC group had lower maximal expiratory pressure (MEP; 94% vs 106% predicted) and reported higher dyspnoea scores (MRC). The PCC group also reported lower levels of physical activity (Frändin/Grimby) and more symptoms consistent with dysfunctional breathing (Nijmegen questionnaire) ($p < 0.001$). Reporting severe symptoms after testing was not associated with abnormal respiratory or functional test results.

Within the PCC group, no sex differences were observed in respiratory or functional outcomes, physical activity level, or dysfunctional breathing symptoms. Women reported higher dyspnoea scores than men ($p = 0.014$).

##### 2.3 Neuropsychological results

**Supplementary table 1. Neurocognitive results**

| | PCC | | Control | | *t* | *p* | Cohen's *d* |
|---|---|---|---|---|---|---|---|
| Test | M | SD | M | SD | | | |
| Ruff 2&7 speed | 46·41 | 11·91 | 52·72 | 8·165 | -2·519 | ·015 | 0·625 |
| CPT-3 detectability | 48·96 | 8·17 | 54·29 | 7·15 | -2·706 | ·009 | 0·701 |
| Coding | 50·29 | 9·93 | 58·8 | 7·97 | -3·961 | <·001 | 0·947 |
| TMT-A | 51·21 | 11·06 | 59·00 | 6·76 | -3·534 | <·001 | 0·856 |
| TMT-B | 41·63 | 16·23 | 54·03 | 11·96 | -3·550 | <·001 | 0·878 |
| CWIT speed composite | 47·07 | 9·47 | 52·13 | 6·18 | -2·602 | ·012 | 0·638 |
| CWIT executive composite | 50·05 | 9·94 | 57·87 | 5·56 | -3·983 | <·001 | 0·982 |
| Digit span composite | 47·01 | 9·68 | 53·38 | 10·35 | -2·655 | ·010 | 0·635 |

Note. All scales were transformed to T-scores. For TMT and CPT-III, scores were reversed so that lower scores indicate poorer performance.
CPT-3= Conners' Continuous Performance Test- 3rd Edition, TMT= Trail Making Test, CWIT= Color-Word Interference Test.

##### 2.4 MRI

The neuroradiological assessment of the conventional MR images yielded Fazekas scores [9] comparable between the groups.

**Supplementary table 2. Fazekas scores.**

| **Fazekas score** | **0** | **1** | **2** | **3** |
|---|---|---|---|---|
| **Patients** | 9 | 22 | 3 | - |
| **Controls** | 9 | 24 | 2 | - |

**Relaxometry**

The relaxometry analysis showed significant differences in white matter volume (p 0.02) and myelin volume (p 0.04), with lower white matter (520 vs 551 ml) and myelin (163 vs 173 ml) volumes in the PCC group compared to controls. There was no significant difference in grey matter volume.

**Diffusion MRI**

The diffusion MRI group comparisons show a significant difference in the right superior-/middle-temporal gyrus ($p < 0.05$, corrected for multiple comparisons across the white matter skeleton) between the PCC group and controls for the scalar metric non-Gaussianity (NG), see Figure S1.

No other diffusion metric (FA, MD, RTOP, MSD, QIV) showed significant differences, although some metrics are close to significance. All metrics trend towards the expected result of impeded white matter integrity with an increase in diffusion and a reduction in anisotropy. The NG group difference does not survive a two-sided test. For conventional dMRI analysis, no significant group differences were found.

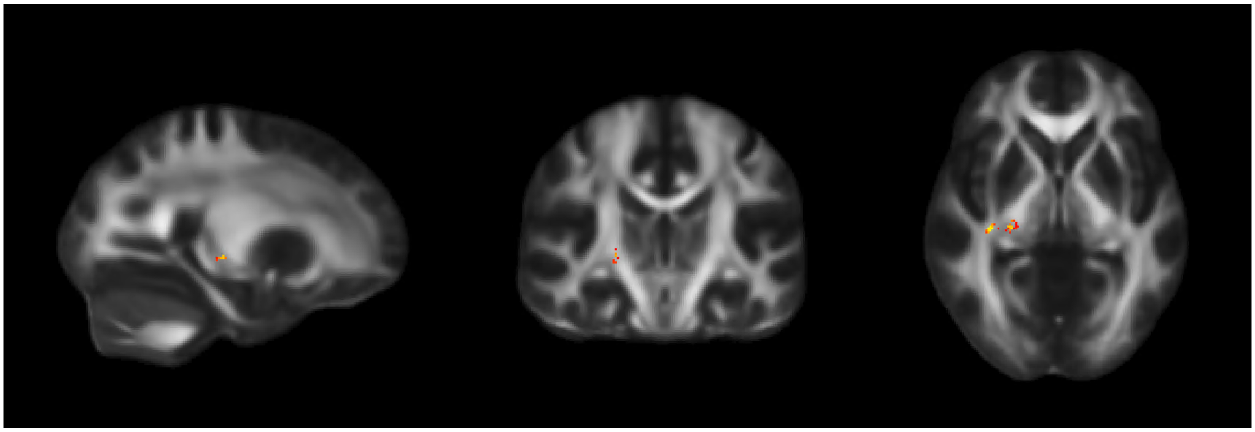

**Figure S1.** Significant group difference in the right superior-/middle-temporal gyrus, showing lower non-Gaussianity in the PCC group compared to controls. N = 69, thresholded at p = 0.05, FWE corrected through TFCE across the white matter skeleton. The lowest corrected p-value is 0.0446.

**Resting-state fMRI**

Resting-state fMRI showed an increase in connectivity for the PCC group in the left central executive, cerebellar, and fatigue related networks compared to controls. See Figures S2, S3, and S4. Group comparisons were also performed for default mode network, salience network, and right central executive network, but these comparisons did not result in any significant differences.

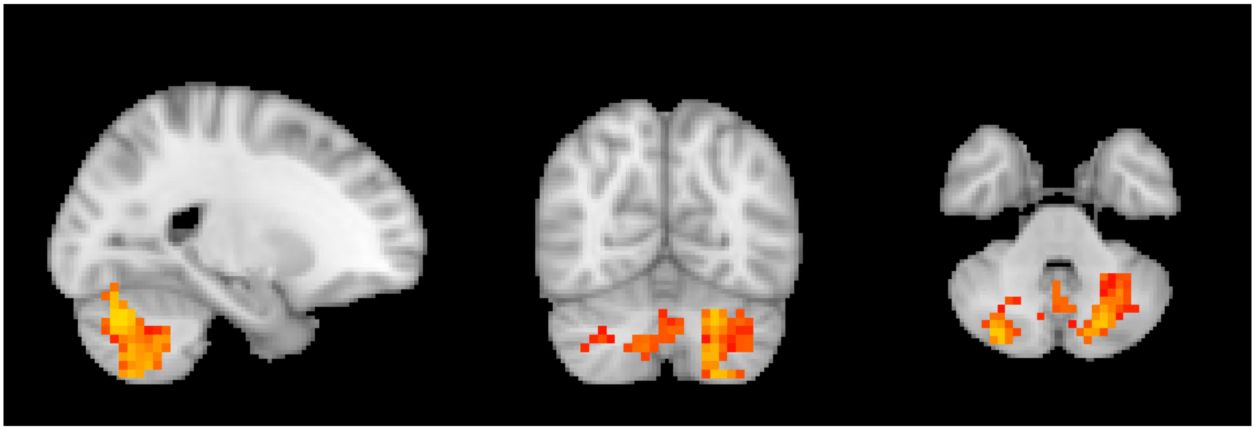

**Figure S2**. Significant group difference in the cerebellar resting state network, showing stronger connectivity in the PCC group compared to controls. N = 69 thresholded at p = 0.05; FWE corrected through TFCE. The lowest corrected p-value is 0.0128. Red indicates that the p-value is close to p = 0.05, while yellow indicates a lower p-value.

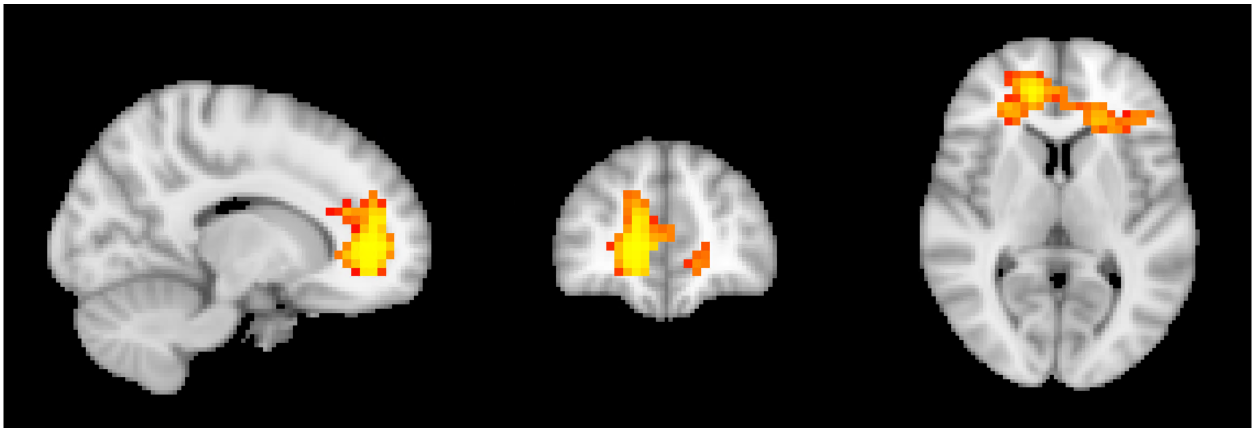

**Figure S3.** Significant group difference in the ventromedial prefrontal cortex and anterior cingulate, interpreted as part of the fatigue resting state network, showing stronger connectivity in the PCC group compared to controls. N = 69 thresholded at p = 0.05, FWE corrected through TFCE. The lowest corrected p-value is 0.0028. Red indicates that the p-value is close to p = 0.05, while yellow indicates a lower p-value.

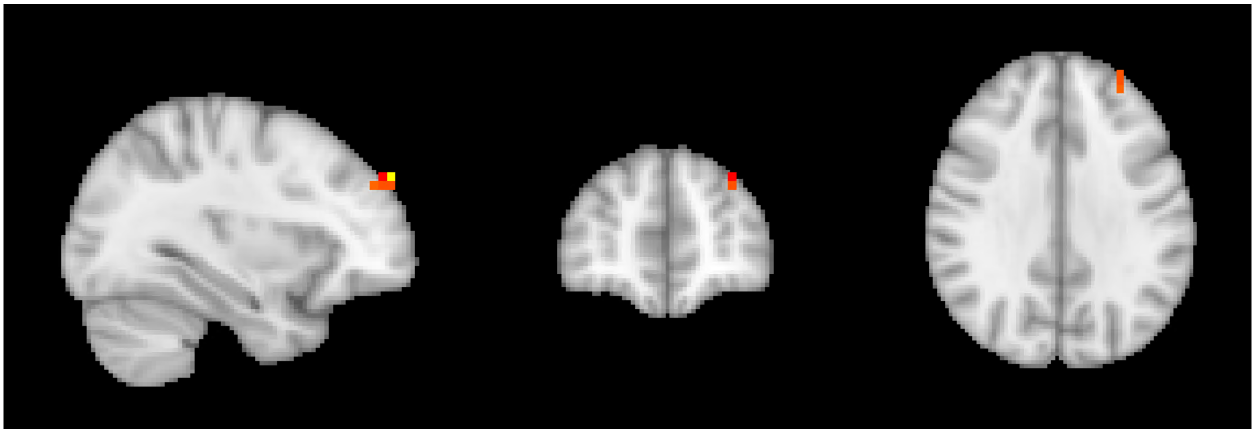

**Figure S4.** Significant group difference in the left central executive resting state network, showing stronger connectivity in PCC compared to controls. N = 69 thresholded at p = 0.05; FWE corrected through TFCE. The lowest corrected p-value is 0.0244. Red indicates that the p-value is close to p = 0.05, while yellow indicates a lower p-value.

##### 2.5 Immunological results

We performed a multiplex immunoassay of 17 inflammatory mediators suggested to be changed by PCC (Supplementary table 3). Although some mediators, including MMP1, MMP8 and the vVWF-A2/ADAMTS13 ratio were significantly changed at the single mediator level, the significance did not remain when taking the multiple comparisons into account. The lack of significant changes most likely reflects a combination of a relatively small material and a lack of strong effects. Given the modest changes observed, it is unlikely that sustained systemic inflammation is a major driver of the symptoms described.

**Supplementary table 3. Cytokines.**

| | **Controls** | | **PCC** | | **P value** | |
|---|---|---|---|---|---|---|
| | **Mean** | **SD** | **Mean** | **SD** | **Raw** | **Corrected** |
| **APP** | 10456 | 1955·0 | 10585 | 2767·2 | 0·414 | n.s. |
| **CCl11** | 256·8 | 31·5 | 265·2 | 45·5 | 0·194 | n.s. |

| | | | | | | |
|---|---|---|---|---|---|---|
| **CCL19** | 149·1 | 124·7 | 207·7 | 430·7 | 0·225 | n.s. |
| **CCL4** | 991·4 | 91·1 | 993 | 136·8 | 0·478 | n.s. |
| **CXCL5** | 575·8 | 193·2 | 671·2 | 356·8 | 0·089 | n.s. |
| **Galectin1** | 38193 | 8962·7 | 40939 | 11320·6 | 0·139 | n.s. |
| **IFNbeta** | 10·08 | 6·1 | 10·32 | 4·2 | 0·426 | n.s. |
| **IL-1beta** | 47·1 | 9·1 | 49·32 | 10·2 | 0·177 | n.s. |
| **IL-1rII** | 8482 | 1636·1 | 8230 | 1386·4 | 0·252 | n.s. |
| **IL-6** | 23·86 | 3·8 | 26·74 | 11·1 | 0·079 | n.s. |
| **IL-7** | 31·12 | 5·2 | 31·73 | 7·0 | 0·345 | n.s. |
| **IL-8 (CXCL8)** | 18·27 | 3·0 | 19·29 | 4·7 | 0·146 | n.s. |
| **MMP1** | 848·9 | 525·1 | 1183 | 828·3 | **0·027** | n.s. |
| **MMP8** | 861·3 | 436·1 | 1429 | 1809·9 | **0·047** | n.s. |
| **P-selectin** | 25985 | 5714·8 | 26262 | 7169·9 | 0·431 | n.s. |
| **TNF** | 26·49 | 4·0 | 25·89 | 6·1 | 0·318 | n.s. |
| **VWF A2** | 2057 | 998·9 | 2234 | 1094·4 | 0·248 | n.s. |
| **ADAMTS13** | 1481616 | 360454·1 | 1346771 | 420391·7 | 0·083 | n.s. |
| **VWF A2/ADAMTS13** | 0·001429 | 0·00071 | 0·001887 | 0·0014 | **0·048** | n.s. |

Note: APP: Amyloid-beta precursor protein, CCL: CC chemokine ligand, IFNbeta: Interferon beta, IL: Interleukin, MMP: matrix metalloproteinase, TNF: tumor necrosis factor, VWF A2: von Willebrand factor A2, ADAMTS: von Willebrand factor-cleaving protease. Comparisons were made with one tailed t-test followed by correction for multiple comparisons. Values are expressed in pg/ml.

##### 2.6 Qualitative Phenomenological Analysis: Two Modal Alterations

Phenomenological philosophy offers accounts of lived experience and embodied subjectivity as being-in-the-world. It is attentive to structures of experience and modal alterations in the subject's very ways of being, experiencing and engaging with others and the world. Phenomenology is consistent with approaches to cognition which understand cognition as "sense-making", emerging from our interaction with the surrounding world. Further, cognitive and affective dimensions of sense-making are understood as intrinsically intertwined, on an experiential level (see for example [41]). This understanding of cognitive and affective dimensions as intertwined also informs the qualitative phenomenological analysis presented in this article. Through the qualitative phenomenological analysis, we identified two modal alterations:

(i) an alteration in which interviewees experienced difficulties to be present in the "here and now", concentrate and "hold" a thought and/or follow a "thread" in thinking and conversations, and correspondingly experienced themselves as "zooming out" and this as entailing an altered sense of self, which we give the title *Difficulties to concentrate and/or follow a "thread" in thinking and conversations;* and

(ii) an alteration in their abilities to execute decisions and actions, which correspondingly entailed an altered sense of themselves, as no longer "themselves" and "slow" or incapable, entitled, below, as *Difficulties to execute decision and actions.*

*(i) Difficulties to concentrate and/or follow a "thread" in thinking and conversations*

The narrated experience of having difficulty to concentrate was commonly combined with experiences of difficulties to "hold" a thought or follow "a thread" in thinking and conversations. As one example, one interviewee explained that "normally, one can follow a thought from the beginning to the end, but I have problem with that." Using metaphorical language, this interviewee explained that "it" – referring to the ability to hold a thought or follow a topic of conversation – "pours away from me, I can't remember what I was talking about" (Int. 3). Another interviewee described how she "lost the thread in conversations," adding that she had to "think a lot because I don't really remember […] I lose details." At times, she said, she "blacked-out completely and could not remember anything" (Int. 14). As yet an example, another interviewee told of how it felt as if she walked "round, round in circles, I can't remember what I should remember […] I need to write down what to do […] I can't just search my memory, I lose it" (Int. 5).[ii]

Further, while experiences such as the above could happen in "calm" situations, they were commonly narrated as part of or worsen in situations with simultaneous stimuli – when too much happened at the same time. Again, some examples can be informative. One interviewee told of how they "got sluggish" or slow in thinking (the Swedish term being used is "*trögtänkt*") when trying to help a friend who was driving the car they were seated in. This interviewee told of how she "couldn't navigate in time, for me this became … shall I ask her to turn here, or should I wait, it became too much, and I felt sluggish," adding that these situations "demand too much of me" (Int. 11).[iii] Not being able to handle such situations, the interviewees said, made them sad. As another example, another interviewee described difficulty with taking in information during work meetings and following conversations with colleagues, and how this also comprised feelings of anger and self-doubt: "Before I started getting help, I was quite angry with myself for not being able to manage it [...] I was very critical of myself... I felt I should be better, you know" (Int. 34).

As can be seen from the excerpts above, at an experiential level, cognitive and affective dimensions are intertwined: the experience of having difficulty to focus or process simultaneous stimuli had affective dimensions (such as sadness), and such affects also shaped how interviewees perceived and experienced themselves (as incapable) and their situations (as overwhelming). In addition, interviewees' narrated experiences of difficulties to concentrate and follow a thread in thinking and conversations also contained experiences of oneself as altered – as 'no longer oneself', as can be illustrated by one participant how stated: "this kind of sluggishness makes me sad, it creates a sense of that this is not who I am" (Int. 11).

*(ii) Difficulties to execute decision and actions*

Interviewees also told of overall alterations in their capability to exercise skills and perform actions in everyday situations. As some examples, one interviewee told of how she, at present, "could not get going with things" (Int. 12), as before. This was also explicitly stated by another interviewee, a health care professional who said, "out of respect for my patients, I don't dare to put myself in such situations [that require fast decisions], as I often don't know how I will react."[v]

Several of the interviewees also narrated detailed experiences of how previously habitual skills no longer functioned at pre-reflective levels. They narrated experiences of how they felt "absent" and "disconnected" (Int. 64) and had to *reflectively attend to how* to engage in everyday activities, in ways that had not previously been the case. They told of a felt lack of ability to be spontaneous, as they "needed to know" and "prepare" to have the energy to execute a plan – be it making a decision or act. As one example, one interviewee told of how she and her husband "were trying to decide what to have for dinner, and I started crying because... it just didn't work, you know. I said: 'I can't do it. I don't know, what do you want from me?'" (Int. 32).

One interviewee – a health care professional – described how they "became sluggish" and had difficulty to perform an everyday task at the hospital, as skills no longer could be performed "automatically." Things "I normally do this fast […] breaking the ampule, taking this needle, it's like I need to go through every step in my head, 'what shall I do', it's like my brain need to recalibrate", this interviewee said, adding that it felt as if "my head can't make these quick decisions" (Int. 11).[vi] In a related way, another interviewee – a chef – told of her experience of having to try to think about what to do next, when chopping onions. This previously embodied skill had become something that needed thematization, i.e. reflective attention to what to do and how to act. This interviewee told of how up to ten seconds could pass when they felt "blank": "I just stand there […] I don't know what to do now," before thinking "mm, fry it!" (Int 16). Further, interviewees narrated how having to think about how to act in mundane situations made them feel that they somehow "couldn't be reached" (In Swedish: "kände mig inte kontaktbar"). This interviewee added that this was the case when "I had to concentrate too much on every little thing I was doing" (Int 26).

At first glance, difficulties in being present in the present – as when zooming out, as in the first theme – and having to explicitly attend to how to act, as in this second theme, may seem like opposites. However, phenomenologically speaking, these can both be seen as instances of a temporary disruption of the "body schema" [41]. The body schema refers to a combination of pre-reflective, practical sensory-motor skills and an implicit, practical awareness of our bodies, motion, and space that make possible the tacit and seamless coordination of different bodily parts in relation to the subject's lived world. It is understood as "the body as it [on a pre-reflective level] actively integrates its positions and responses in the environment" [42(p.548)]. As shown in past research, illness commonly disrupts the body schema when it makes the body manifest as an obtrusive or painful object one needs to attend to, which disrupts the embodied self's previously seamless engagement with the world[43,44]. In the present case, however, the zooming out *and* the needing to thematically attend to how to act, breaking down an action into smaller steps, bring out a distinct combination of what can be referred to as a *slackening of one's directedness* within a particular situation, where this slackening, i.e., this temporary loss of focus and orientation, seems to be part of the *forced* thematization of action where explicit attention is required to perform things one previously did not need to think about (hence, having to break down one action into smaller segments, attending to each of them, for it to be possible to perform the action).

What these interviewees otherwise just would do – cook or prepare a needle – now requires attention and effort: it required work, and patience, as things took time when mundane activities no longer could be pre-reflectively and tacitly performed. This was narrated as frustrating, stressful, sad, and embarrassing. Further, interviewees said, when this happened – when they had to think about, and thematize everyday small parts of an action –, they sometimes lost their temper (Int 16, 3, 51) and then felt bad about losing their temper. As part of these experiences, interviewees told of an altered sense of self – how they felt slow and incapable, and different from before.

*Intertwinement of (i) and (ii)*

The examples of the modal alterations (i) and (ii) were narrated as distinct experiences in the interviews, and half of the interviewees who told of what we have identified as the first kind of interview also told of the second kind of experience. Further, these experiences were also, at times, narrated as temporally intertwined, one leading to the other. As but one such example, one interviewee told of the experience of spatiality as intertwined with difficulties in making sense of what happened around them, and acting appropriately, when, again, much happened at the same time. This interviewee told of how she experienced an everyday task of taking a bus as a "horrifying experience:" everything around her, she said, "moved so fast" and "[when] we moved at five kilometers per hour, it felt like 120" (Int 30). Such spatial experiences, combined with for example sensitivity to sound, were part of an experience of difficulty to take in what happened around her, fast enough, which also fed into the felt difficulty to execute a simple task: "Then it was like I could barely remember that I needed to press the button to get off, you know... you have to press to indicate you want to get off, and I just forgot" (Ibid).[iv] This inability led to feelings of loneliness and isolation from the world.

As a general feature, across (i) and (ii), interviewees narrated distinct losses: one could no longer engage with others and the world as before, which was combined with a sense of oneself as no longer "oneself". Such loss was also combined with a basic doubt not only in one's own abilities, but also in whether one would ever get back to one's previously taken-for-granted ways of being, experiencing and engaging with others and the world, and this as felt as existentially challenging. Phenomenologically, this can be understood as exemplifying what Havi Carel [46(p.184)] has termed a basic "bodily doubt." In Carel's analysis, the basic bodily doubt is part of "existential feelings," i.e., pre-intentional affective "background orientations" that shape the self's felt relation to the world, and the very ways in which both self and world are – and can be – experienced [46 (p.46),47]. The basic bodily doubt is not only cognitive in the sense of a "disruption of belief," but entails a *foundational disruption of one's sense of trust, certainty and confidence in one's own body and bodily capacities* [46(p.184)]. Further, it can "descend at any moment", thus implying a thorough unpredictability that makes it particularly threatening and difficult to incorporate into one's everyday lifeworld, and it "invades the normal sense" of the everyday, resulting in a feeling of exposure and threat [46(p.185)]. This resonates with the interviewees' narrated lived experience, where the experienced difficulties to concentrate, follow a "thread" in thinking and conversation, and in executing decisions and actions were narrated as feeding into everyday life, in a way that was felt as existentially challenging.

Further, a basic bodily doubt, as a dimension of existential feelings can be part of many different illness experiences, but the present analysis brings out a specific case. The basic bodily doubt, here, seems to be centred on and about the self's own cognitive faculties – i.e., *about faculties that are deeply involved in the everyday tacit assessing of one's capacities to engage with the others and the world*. This is a doubt about previously taken-for-granted bodily, cognitive capacities, and it reveals a basic vulnerability, as basic assumptions of predictability are overthrown. It shapes and constrains what is felt as possible.

#### 2.7 Correlation analysis

Correlation analysis of MRI and neuropsychology was performed for controls and PCC separately, for combinations of networks and psychometric scores which showed significant interactions.

DAN connectivity and CWIT executive; the significant interaction effect shown in Figure 2 in the main manuscript seems to be explained by the fact that controls have significant negative correlation between DAN and CWIT executive composite score in the ventral part of the intraparietal sulcus, while the PCC group have significant positive correlation between DAN and CWIT executive composite score in the intraparietal sulcus closer to vertex. See Figures S5 and S6.

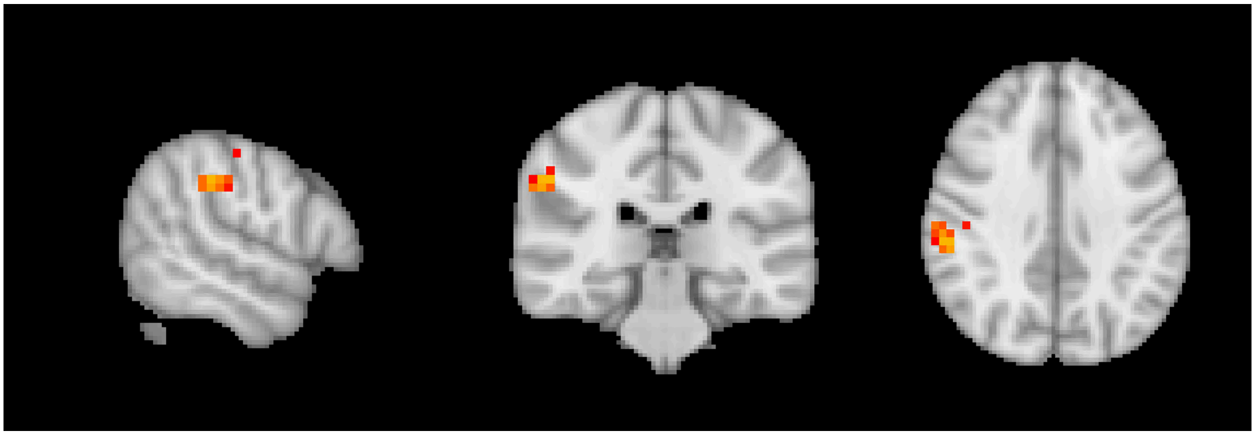

**Figure S5.** Significant negative correlation between DAN and CWIT executive composite score for controls, N = 34, thresholded at p = 0.05, FWE corrected through TFCE. The lowest corrected p-value is 0.013. The PCC group has no significant negative correlations between DAN and CWIT. Red indicates that the p-value is close to p = 0.05, while yellow indicates a lower p-value.

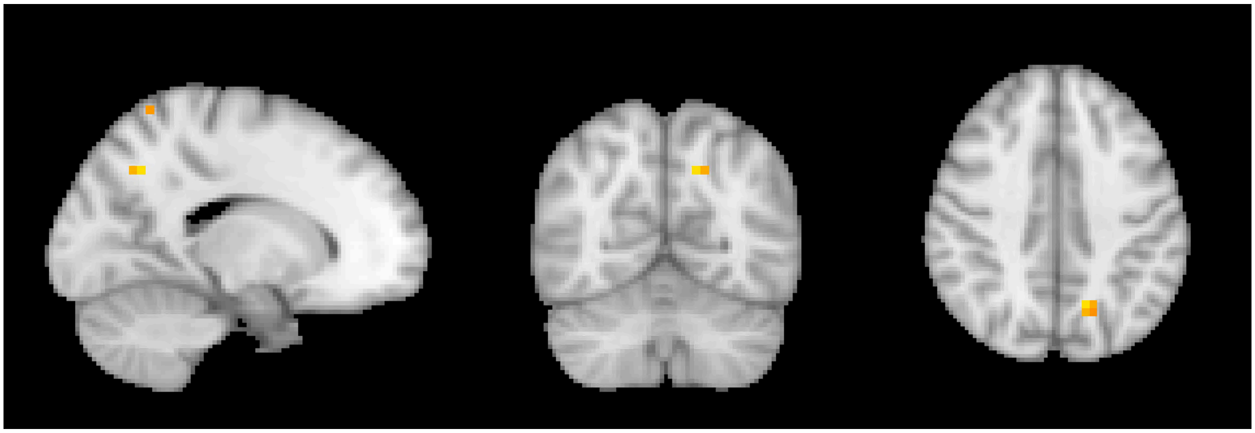

**Figure S6.** Significant positive correlation between DAN and CWIT executive composite score for the PCC group, N = 31, thresholded at p = 0.05; FWE corrected through TFCE. The lowest corrected p-value is 0.006. The control group has no significant positive correlations between DAN and CWIT executive composite score. Red indicates that the p-value is close to p = 0.05, while yellow indicates a lower p-value.

Salience connectivity and coding; the significant interaction effect shown in Figure 3 in the main manuscript seems to be explained by the fact that the PCC group has a negative correlation between salience and the Coding subtest in the insula, while the controls lack this correlation. See Figure S7.

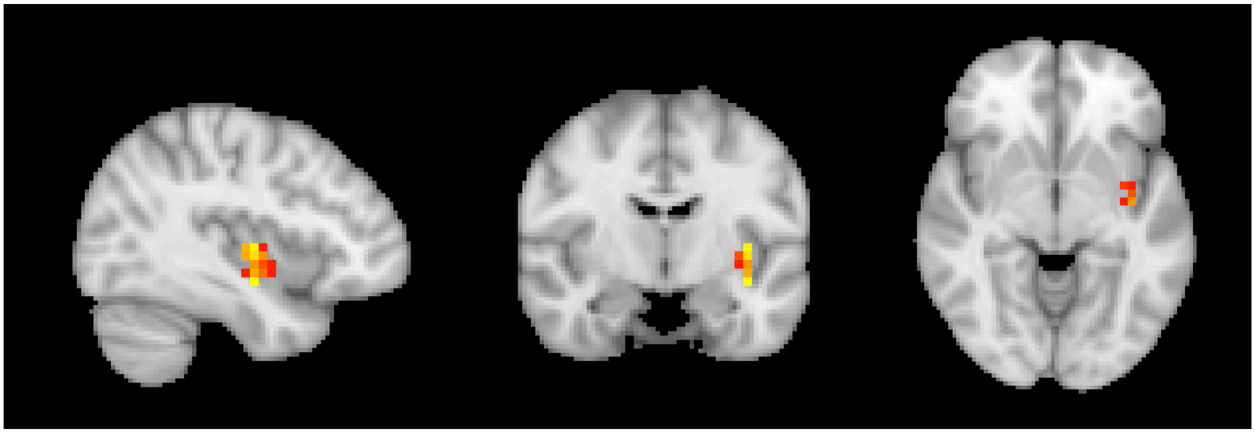

**Figure S7.** Negative correlation between salience and coding for the PCC group, N = 31, thresholded at p = 0.1, FWE corrected through TFCE. The lowest corrected p-value is 0.068. For the controls, the lowest corrected p-value is 0.674. Red indicates that the p-value is close to p = 0.1, while yellow indicates a lower p-value.

Left executive network and CWIT; the significant interaction effect shown in Figure 4 in the main manuscript seems to be explained by the fact that controls have negative correlation between left executive network and CWIT executive composite score, while the PCC group lack this correlation. See Figure S8. The PCC group instead has a significant positive correlation in the right cerebellum (p = 0.0154).

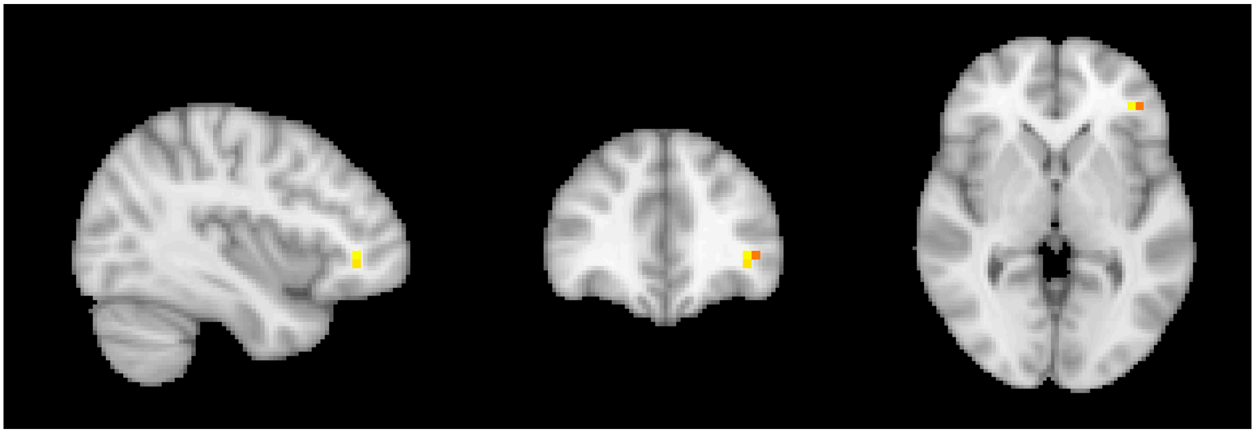

**Figure S8.** Negative correlation between left executive network and CWIT executive composite score for controls, N = 34, thresholded at p = 0.2, FWE corrected through TFCE. The lowest corrected p-value is 0.107. For the PCC group, the lowest corrected p-value is 0.357. Red indicates that the p-value is close to p = 0.2, while yellow indicates a lower p-value.

**Diffusion MRI and immunology**

The significant interaction effect between RTOP and MMP1 (Figure 5) seems to be explained by the fact that controls have a widespread positive correlation between RTOP and MMP1, while the PCC group instead has a widespread negative correlation. See Figures S9 and S10.

Furthermore, the PCC group has a significant correlation between RTOP and WFA2 (lowest corrected p-value 0.001) and between RTOP and MMP8 (lowest corrected p-value 0.0042), not shown. The controls do not have any significant correlations between RTOP and WFA2 or MMP8, but the difference is not significant (no interaction).

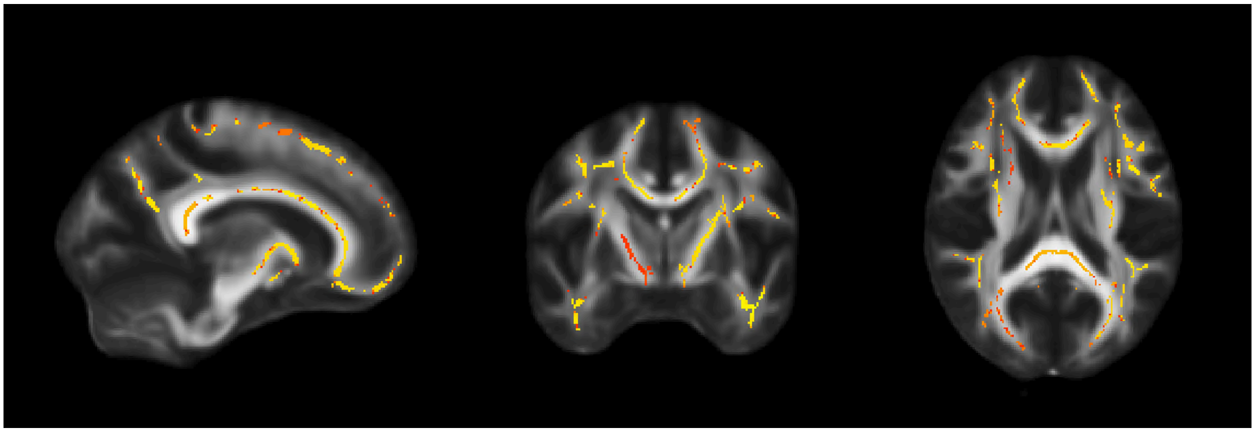

Figure S9. Positive correlation between RTOP and MMP1 for controls, N = 28, thresholded at p = 0.2, FWE corrected through TFCE. The lowest corrected p-value is 0.1042. For the PCC group, the lowest corrected p-value is 0.867. Red indicates that the p-value is close to p = 0.2, while yellow indicates a lower p-value.

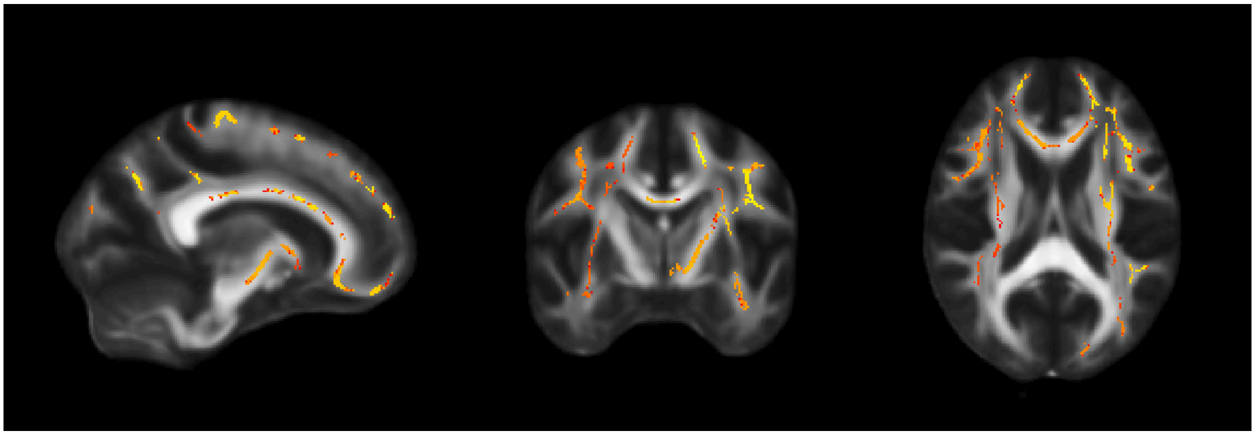

Figure S10. Negative correlation between RTOP and MMP1 for the PCC group, N = 25, thresholded at p = 0.2, FWE corrected through TFCE. The lowest corrected p-value is 0.0508. For controls, the lowest corrected p-value is 0.937. Red indicates that the p-value is close to p = 0.2, while yellow indicates a lower p-value.

The significant interaction between MD and WFA2 (Figure 6) seems to be explained by the fact that the PCC group has a significant negative correlation between MD and WFA2 (lowest corrected p-value 0.0022), while it is not significant for the controls. Furthermore, the PCC group has a significant negative correlation between MD and MMP8 (lowest corrected p-value 0.0114). See Figures S11 and S12. The controls do not have any significant correlation between MD and MMP8, but the difference is not significant (no interaction).

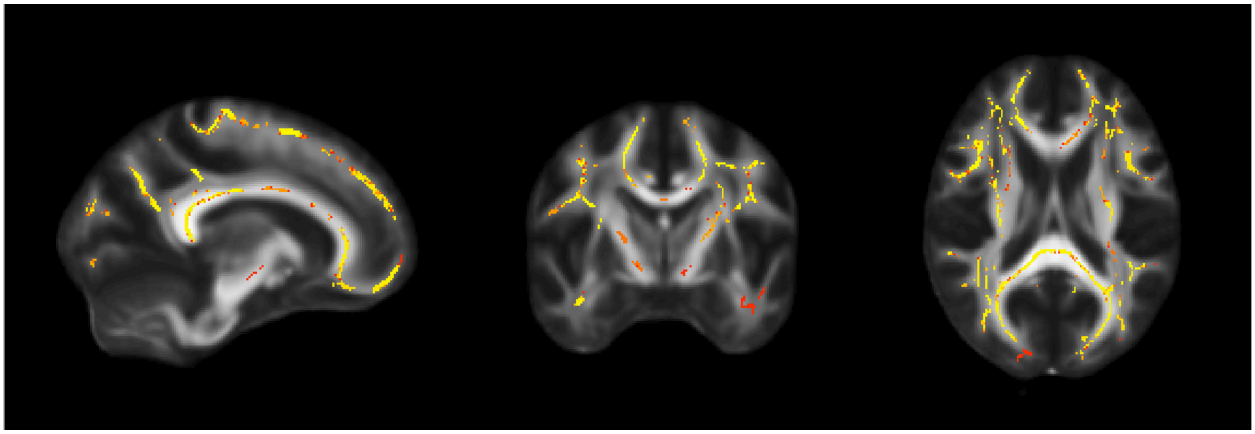

**Figure S11.** Significant negative correlation between MD and WFA2 for the PCC group, N = 25, thresholded at p = 0.05, FWE corrected through TFCE. The lowest corrected p-value is 0.0022. For controls, the lowest corrected p-value is 0.2054. Red indicates that the p-value is close to p = 0.05, while yellow indicates a lower p-value.

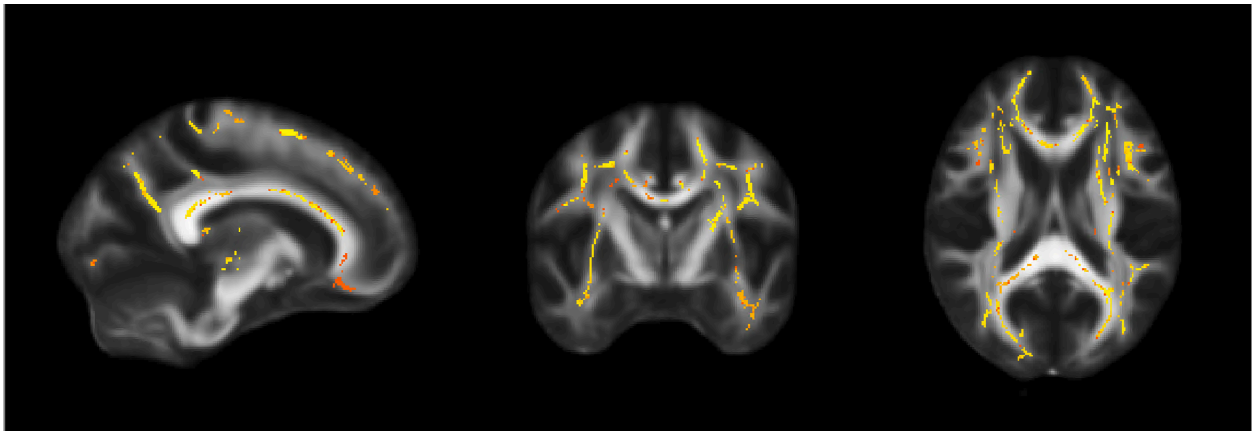

**Figure S12.** Significant negative correlation between MD and MMP8 for the PCC group, N = 25, thresholded at p = 0.05, FWE corrected through TFCE. The lowest corrected p-value is 0.0114. For controls, the lowest corrected p-value is 0.3008. Red indicates that the p-value is close to p = 0.05, while yellow indicates a lower p-value.

Furthermore, the PCC group has a significant positive correlation between FA and WFA2, and between FA and MMP8. The controls do not have any significant correlations, but the difference is not significant (no interaction).

The PCC group has a significant negative correlation between MSD and WFA2, and between MSD and MMP8. The controls do not have any significant correlations, but the difference is not significant (no interaction).

The PCC group has a significant negative correlation between QIV and WFA2, and between QIV and MMP8. The controls do not have any significant correlations, but the difference is not significant (no interaction).
